# Megabase-scale human genome rearrangement with programmable bridge recombinases

**DOI:** 10.1101/2025.05.14.653916

**Authors:** Nicholas T. Perry, Liam J. Bartie, Dhruva Katrekar, Gabriel A. Gonzalez, Matthew G. Durrant, James J. Pai, Alison Fanton, Masahiro Hiraizumi, Chiara Ricci-Tam, Hiroshi Nishimasu, Silvana Konermann, Patrick D. Hsu

## Abstract

Bridge recombinases are a class of naturally occurring RNA-guided DNA recombinases. We previously demonstrated they can programmably insert, excise, and invert DNA *in vitro* and in bacteria. Here, we report the discovery and engineering of IS622, a simple two-component system capable of universal DNA rearrangements of the human genome. We define strategies for the optimal application of bridge systems, leveraging mechanistic insights to improve their targeting specificity. Through rational engineering of the IS622 bridge RNA and deep mutational scanning of its recombinase, we achieve up to 20% insertion efficiency into the human genome and genome-wide specificity as high as 82%. We further demonstrate intra-chromosomal inversion and excision, mobilizing up to 0.93 megabases of DNA. Finally, we provide proof-of-concept for excision of a gene regulatory region or expanded repeats relevant for the treatment of genetic diseases.

## INTRODUCTION

Genomes are the foundational information layer that encode biological complexity, from the activity of individual enzymes to coordinated cellular networks that orchestrate organism-level behavior (Hartwell et al. 1999; Jacob & Monod 1961). These genotype-to-phenotype relationships are determined by combinations of mutations across megabase-scale genomic regions, limiting our understanding of even the simplest genomes. The development of RNA-guided technologies such as CRISPR genome engineering has revolutionized our ability to interrogate biological function through programmable DNA modification (Hsu et al. 2014; Pacesa et al. 2024). However, these approaches rely on complex multi-step mechanisms such as homology-directed repair or prime editing that restrict the length scale of possible genome edits (Anzalone et al. 2019; Liao et al. 2024). In contrast to CRISPR, DNA recombinases such as Cre can mobilize many thousands of DNA base pairs, but require pre-installed recognition sites to enable mammalian genome recombination and leave residual scars (Koeppel et al. 2024; Pinglay et al. 2025; Sauer 1987; Sauer & Henderson 1988). While fusion to RNA-guided systems or directed evolution can modify the sequence specificity of recombinase enzymes, such tools are largely limited to gene integration and leave long insertion site scars (Anzalone et al. 2021; Fanton et al. 2024; Pallarès-Masmitjà et al. 2021; Yarnall et al. 2022).

As the first single-effector class of RNA-guided DNA recombinases, bridge recombinases have the potential to combine the flexible specificity of CRISPR with the payload scale of recombinases (Durrant et al. 2024). Derived from IS110 insertion sequence elements, bridge recombinases recognize their target and donor DNA using a bridge RNA (bRNA) that confers programmable specificity for both substrates, enabling a universal mechanism for DNA insertion, inversion, and excision (**Figure 1A**). In contrast, CRISPR-associated transposases specify only the target DNA via a guide RNA, while requiring a fixed DNA donor sequence that limits them to gene insertion (Klompe et al. 2019; Lampe et al. 2023; Strecker et al. 2019; Tou et al. 2023). Structural studies of the bridge recombinase IS621 revealed that two recombinase dimers bind both bRNA loops separately and then bind the cognate DNA to form a tetrameric synaptic complex (Hiraizumi et al. 2024) (**Figure 1B**). Strand exchange occurs after the formation of covalent bonds between the DNA and the recombinase, resulting in a Holliday junction intermediate that is resolved to yield the final recombination product (**Figure 1C**).

**Figure 1:**
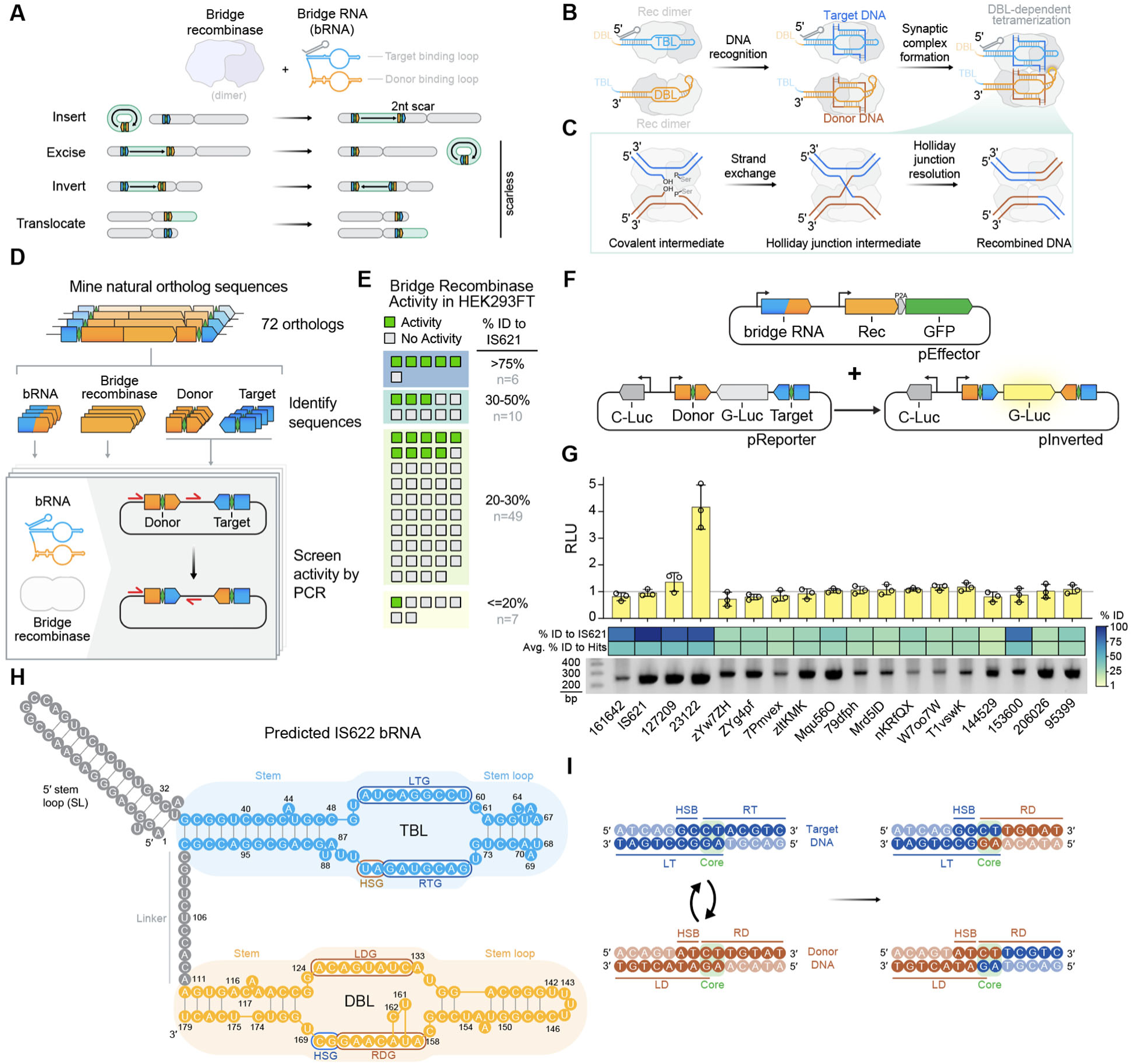
Activity screening of bridge recombinases in human cells (**A**) Schematic of chromosome-scale genome modifications with bridge recombination. (**B**) Overview of bridge recombinase synaptic complex assembly, based on IS621. TBL, target-binding loop; DBL, donor-binding loop. (**C**) Overview of strand exchange, Holliday junction formation, and resolution within a bridge recombinase transpososome, based on IS621. (**D**) Schematic of ortholog screening approach. Key components of bridge recombinase systems from IS110 elements were encoded on reporter plasmids, where inversion between target and donor is measured to determine activity. (**E**) Activity of bridge recombinases based on PCR detection of the inversion product junction. Orthologs are categorized by percent identity to IS621. (**F**) Schematic of C-luciferase/G-luciferase inversion reporter assay. (**G**) Activity of orthologs in luciferase assay. Amplification of inversion junction shown below. Percent identity to IS621 and percent identity to other active orthologs is shown. RLU, relative luminescence units. (**H**) Schematic of IS622 bRNA, based on IS621 bRNA structure (PDB: 8WT6). LTG, left target guide; RTG, right target guide; LDG, left donor guide; RDG, right donor guide; TBL, target-binding loop; DBL, donor-binding loop; HSG, handshake guide. (**I**) Target and donor DNA recognized by the IS622 bRNA. Recombination occurs between CT cores. LT, left target; RT, right target; LD, left donor; RD, right donor; HSB, handshake bases.

Our initial study provided proof-of-concept for using bridge recombinases in programmable modification of prokaryotic genomes (Durrant et al. 2024). Here, we describe the discovery of a bridge recombinase with high activity in human cells, termed IS622. We enhance its activity by rational engineering of its bRNA via point mutations and scaffold modifications. Using these enhanced bRNAs, we discovered design principles for maximizing the specificity of insertion into the human genome, achieving as high as 82% specificity. To further augment activity of IS622, we performed a deep mutational scan of the recombinase directly in human cells. Finally, we combine a rationally engineered high activity recombinase mutant with our enhanced bridge RNAs, achieving up to 20% insertion efficiency into the human genome. Beyond gene insertion, we use IS622 for programmable, precise, and scarless genome rearrangements, inverting up to 0.93 Mb and excising up to 0.13 Mb with no apparent distance dependency. Finally, we demonstrate therapeutic proof-of-concept with excision of the *BCL11A* enhancer for sickle cell anemia and of repeat sequences found in Friedreich’s ataxia.

## RESULTS

### Diverse bridge recombinases are active in human cells

We previously characterized reprogrammable bridge recombination *in vitro* and in *E. coli*. In our previous efforts to effectively use prokaryotic enzymes in eukaryotic systems, broad screening of diverse orthologs with a wide range of activity was necessary before an optimal ortholog was selected for further development (Durrant et al. 2022; Konermann et al. 2018). Leveraging our database of IS110 elements (Durrant et al. 2024), we predicted the recombinase coding sequences and cognate bRNAs for 72 diverse IS110 insertion sequence elements (**Figure S1A-C**). Using the IS110 element boundaries determined by comparative genomics, we reconstituted the natural target and donor sequences and tested them in an inversion reporter assay in human cells (**Figure 1D**). The inverted target/donor junction was detected for 18/72 (25%) orthologs, confirming that at least some bridge recombinases do not require prokaryotic host factors to function in human cells (**Figure 1E**).

Next, we quantified the efficiency of these 18 functional orthologs by measuring inversion of Gaussia luciferase (**Figure 1F**). In this assay, only a single ortholog exhibited activity above the limit of detection, despite all orthologs having observable activity via amplification of the inversion junction (**Figure 1G**). The active ortholog, previously labeled 23122 (Durrant et al. 2024), with a 4-fold change over background, is from *Citrobacter rodentium*, and identified as ISCro4 in ISFinder (Siguier et al. 2006). We named this ortholog IS622, given its similarity to the previously described IS621 ortholog (88% recombinase and 86.6% bRNA identity) (**Figure S2A-B**). A comparison of an AlphaFold3-predicted model of the IS622 recombinase with our cryo-EM structure of the IS621 recombinase (Hiraizumi et al. 2024) confirmed the high structural similarity between the recombinases, with root mean square deviation (RMSD) of 2.5 Å for equivalent Cα atoms **(Figure S3A-D)**, allowing modeling of the likely bRNA structure (**Figure 1H**).

The model suggests that the IS622-bRNA complex recognizes 14 nucleobases in both of its donor and target DNA substrates, using its target- and donor-binding loops (TBL and DBL) in the bRNA, respectively. These two bRNA loops recognize both strands of the target and donor sequences, resulting in conservative recombination around a dinucleotide core sequence (**Figure 1I**). Additionally, both loops encode handshake guides (HSG) that base-pair upstream of the core sequences with the handshake bases (HSB) after top strand exchange, a mechanism that drives the recombination reaction forward (Hiraizumi et al. 2024) (**Figure S4**).

### Structure-guided bridge RNA engineering

We next reasoned that rational engineering of the bRNA may yield increased activity of IS622 in human cells, as guide RNA engineering has been a longstanding strategy for increasing the activity of CRISPR systems in human cells (Cong et al. 2013; Hsu et al. 2013; Kim et al. 2021). Despite encoding specificity for both DNA substrates on a single RNA molecule, our structural study of IS621 suggested that two distinct bridge RNA molecules may be employed during recombination by the tetrameric complex (Hiraizumi et al. 2024). Building on this observation, we asked if productive transposome assembly in human cells would occur if the TBL and DBL were expressed separately (**Figure 2A**). Using an mCherry inversion reporter, we compared the full length bRNA with two versions of a split bRNA, with or without the 5′ stem loop. We observed that split bRNAs were not only functional, but resulted in ∼2.1-fold increase in recombination efficiency by mCherry mean fluorescence intensity (MFI) (**Figure 2B-C**).

**Figure 2:**
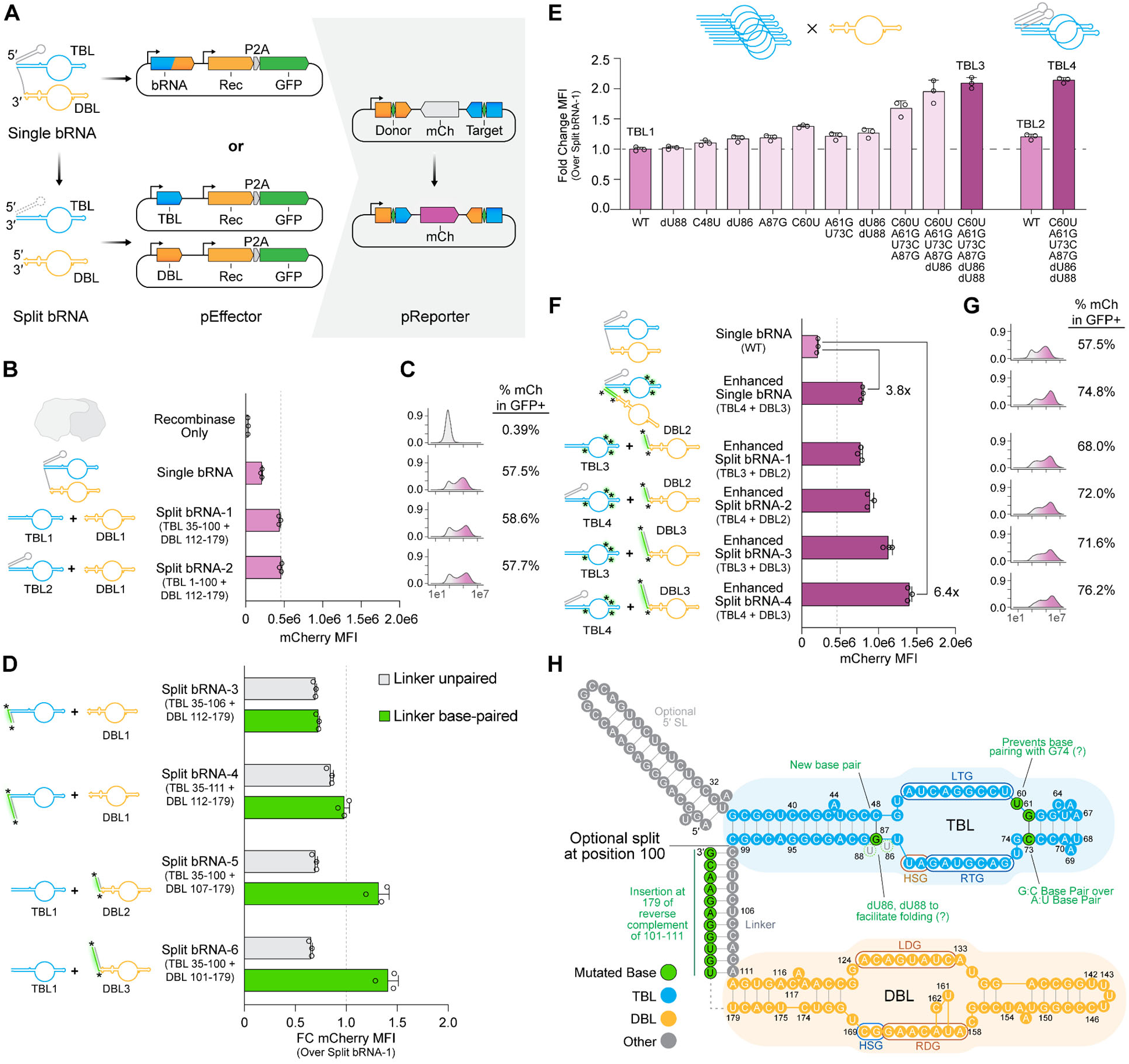
Bridge RNA engineering for efficient recombination **(A**) Overview of recombinase and bRNA expression plasmids with mCherry inversion reporter. (**B**) Inversion efficiency with single and split bridge RNAs. TBL and DBL intervals based on 179nt single bridge RNA. MFI, mean fluorescence intensity. TBL, target binding loop; DBL, donor binding loop. (**C**) Inversion efficiency as measured by percentage of mCherry+ cells within GFP+ (recombinase expressing) cells. (**D**) Inversion efficiency (MFI) of TBL and DBL with extended stems. All or part of the bRNA linker sequence (positions 101-111) was included on a TBL or DBL, with or without insertion of a complementary sequence. Represented as fold-change compared to Split bRNA-1 in (B). (**E**) Inversion efficiency of mutated TBLs when paired with DBL1. TBL2 and TBL4 include the 5′ stem loop of bridge RNA (positions 1-34). (**F**) Inversion efficiency (MFI) of single and split bRNAs featuring point mutations and extended stems. (**G**) Inversion efficiency (% mCherry+) of single and split bRNAs featuring point mutations and extended stems. (**H**) Schematic of enhanced bRNA. Green text indicates enhancing mutations relative to the WT bRNA.

In the WT bRNA, the TBL and DBL are connected via a linker region that connects the stem regions of the two loops to one another. When determining the optimal region to split the bRNA into its two loops, we found that inclusion of part or all of the linker region (positions 101-111) onto either loop significantly reduced activity relative to eliminating the linker (**Figure 2D**, grey boxes). While the TBL contains a 12 bp continuous stem with a single base (A44) flipped out, the DBL stem contains three unpaired positions in the middle flanked by 3 and 4 paired bases on each side (**Figure 1H**). We therefore asked if increased stem length would modulate activity. By inserting bases to pair with the natural unpaired linker region, we increased the TBL stem length by 6 bp or 11 bp. However, we did not observe an increase of activity compared to Split bRNA-1, which terminates the TBL at the end of its natural stem. In comparison, lengthening the shorter DBL stem by 6 bp (DBL2) and 11 bp (DBL3) resulted in 1.3-fold and 1.4-fold greater activity over DBL1, respectively (**Figure 2D**, green boxes). Taken together, we find that limiting unpaired linker bases and modulating stem length are effective at increasing recombination activity in a loop-specific manner.

We next tested the effects of rational substitutions and deletions in both loops, focusing on maximizing the number of base paired nucleotides to help stabilize the bRNA structure. We initially assessed these changes on each loop separately using the split bRNA architecture. While DBL mutations yielded limited improvement over the unmodified split system (0.2 to 1.1-fold change) (**Figure S5**), substitutions and deletions in the TBL were more successful (1.0 to 1.4-fold change) (**Figure 2E**). We next combined these three substitutions and two deletions with an A:U→G:C base-pair modification (A61G + U73C) and observed cumulative increases in activity, culminating in TBL3 (2.1-fold activity over TBL1). Inclusion of the natural 5′ stem loop (TBL4) additionally slightly increased activity (2.1-fold over TBL1), similar to our previous study of the IS621 complex (Hiraizumi et al. 2024).

We next modified the single bRNA scaffold to contain TBL4 and DBL3, which each performed optimally in the split system. We observed a 3.8-fold increase in activity in the context of the single bRNA (**Figure 2F-G**) and named this the enhanced bRNA. To determine the optimal split bRNA configuration, we tested all four combinations of the top 2 TBLs and top 2 DBLs. The best split loop pairing, TBL4 with DBL3, exhibited a 6.4-fold increase over the WT bRNA, highlighting synergistic effects of individual loop modifications.

Modeling of our enhanced bRNA supports potential mechanistic explanations for the improved activity (**Figure 2H**). Multiple mutations (A87G, A61G + U73C) in the TBL that were observed to increase activity likely contribute to stabilizing the TBL stem by replacing two existing base pairs with a stronger G:C pair, while C60U (the strongest individual mutation) is expected to help delineate the TBL stem boundary by preventing undesirable baseparing with G74 (**Figure 2E**). We are able to find that the unpaired U86 and U88 bases on the other hand were dispensable, and that their removal in fact enhanced activity (dU86 and dU88) (**Figure 2E**). The linker region between the TBL is expected to not be in contact with the recombinase, and seems to be tolerant of being base-paired via extension of the DBL stem, likely contributing to its additional stabilization. Finally, in contrast to previous reports on engineering a Cas12f guide RNA (Kim et al. 2021), modifying poly-U sequences in the DBL did not increase activity by increasing expression from a U6 promoter (**Figure S5**).

### DBL-DBL recombination with IS622

The 14 nt target sequence naturally recognized by the IS622 bRNA appears only once in the human genome, making it a straightforward test case for sequence-specific insertion. We co-transfected the recombinase and WT bRNA into human cells with a plasmid encoding the WT donor sequence and a constitutive puromycin marker (**Figure 3A**). After 18 days of selection to enrich for both on- and off-target insertions, we measured the relative efficiency of insertion genome-wide via deep sequencing (**Methods**). While we observed insertion reads mapping to the expected target, these represented only 0.05% of insertions. Surprisingly, the consensus insertion sequence across all integration sites closely matched the donor sequence, rather than the target sequence (**Figure 3B**). Further inspection of off-target sites showed that 91.3% of insertions were more similar to the plasmid donor sequence than the expected genomic target (**Figure 3C**). Out of all insertions, 75.5% were within 4 nt Levenshtein distance from the WT donor sequence, including 1.59% into human genome occurrences of the WT donor sequence. In contrast, only 0.25% of insertions were within 4 nucleotides of the WT target sequence (**Figure S6A-B**). The higher similarity of off-targets to the sequence specified by the DBL suggested a distinct sequence recognition mechanism for IS622.

**Figure 3:**
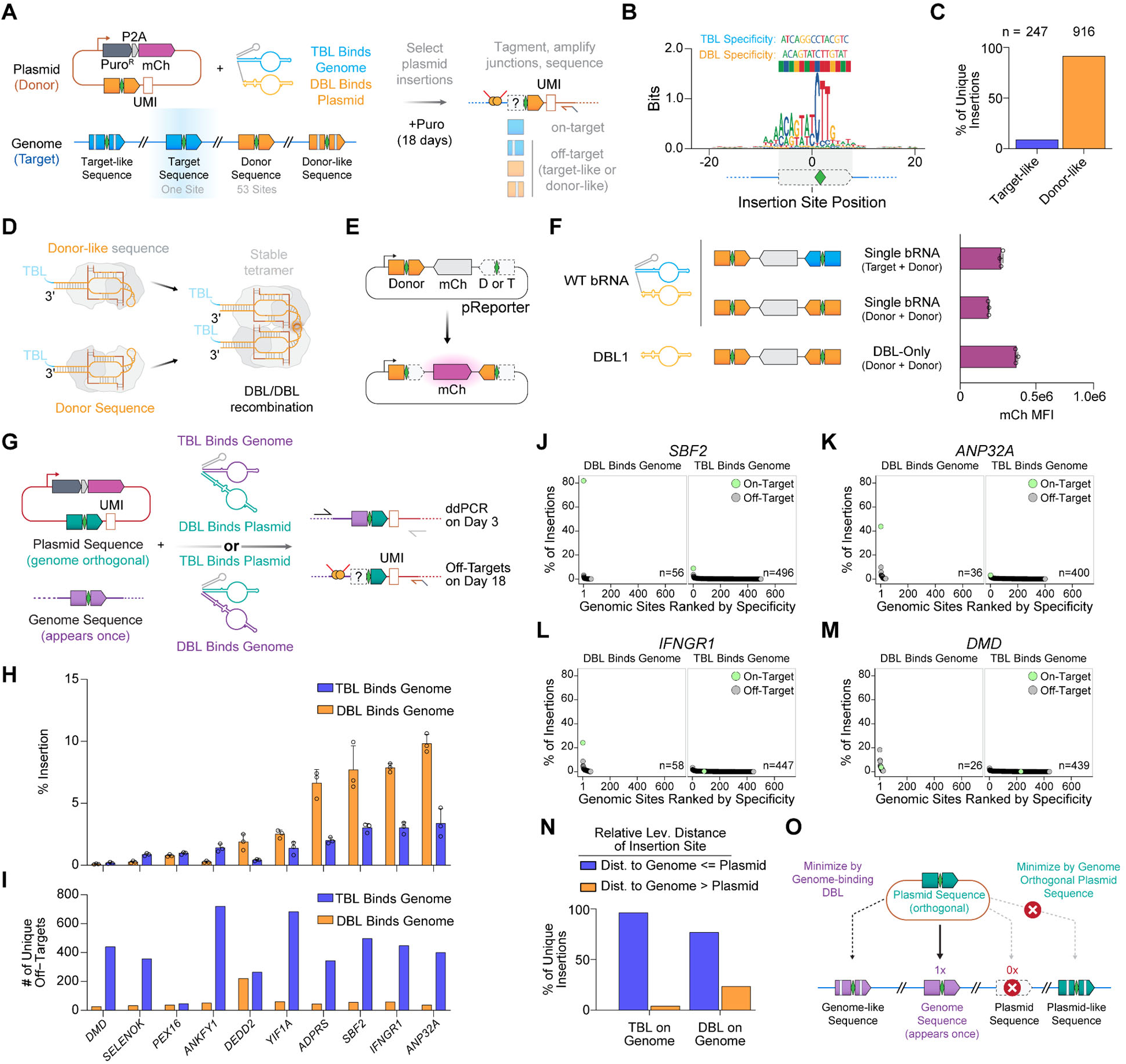
DBL binding to the genome enables high-specificity insertion (**A**) Schematic of genome-wide insertion specificity assay, with WT bRNA shown as an example. On- and off-targets are measured after insertion of puromycin resistance gene and antibiotic selection. UMIs are counted genome-wide to determine specificity across all insertions. UMI, unique molecular identifier. (**B**) Sequence logo of all insertion sites using the WT bRNA, weighted by observed UMIs. TBL and DBL recognition sequences are shown. (**C**) Percentage of all insertions binned by similarity to IS622 WT target sequence or WT donor sequence. (**D**) Schematic of DBL-DBL mediated recombination. DBL RNPs bind DBL-like DNA and tetramerize, yielding recombination. (**E**) Schematic of mCherry inversion reporter. (**F**) Inversion efficiency of split and single bRNAs for target+donor or donor+donor recombination. MFI, mean fluorescence intensity. (**G**) Schematic of approach for measuring efficiency and specificity of insertion into the genome sites using both enhanced bRNA orientations. (**H**) Insertion efficiency by digital droplet PCR three days post-transfection (no selection) at 10 genomic loci using both bRNA orientations. (**I**) Number of unique off-targets observed for each locus with each bRNA orientation. (**J**) Insertion specificity of *SBF2* using both bRNA orientations after 18 days of selection. (**K**) Insertion specificity of *ANP32A.* (**L**) Insertion specificity of *IFNGR1.* (**M**) Insertion specificity of *DMD.* (**N**) Percentage of unique insertions binned by the relative Levenshtein distance of the insertion site to the intended genome site or the plasmid encoded sequence. Results cumulative of all 10 loci in (I). (**O**) Model for minimizing off-target insertions with IS622 bridge recombinases.

In our previous study of insertion specificity of IS621 in *E. coli*, we had observed low frequency recombination between the plasmid-encoded donor sequence and donor-like sequences matching the DBL in the genome (Durrant et al. 2024). Structural and biochemical studies indicated that this “donor-donor” reaction facilitated by IS621 is likely a result of transpososomes containing two DBLs from separate bridge RNAs, rather than one TBL and one DBL (**Figure 3D**) (Hiraizumi et al. 2024). DBL-DBL, but not TBL-TBL, transpososomes are possible due to the unique stem-loop of the DBL, which is involved in tetramer stabilization (Hiraizumi et al. 2024) (**Figure S2B, Figure S6C-D**).

Given that donor-like insertion sites (matching the DBL instead of the TBL) were much more abundant, we employed an mCherry inversion reporter to compare recombination between target-donor and donor-donor with the WT bRNA, confirming that both recombination types can occur robustly (**Figure 3E-F**). We then delivered only a DBL from the split design, demonstrating that it is sufficient for recombination and in fact leads to higher donor-donor recombination than the full-length single bridge RNA containing both DBL and TBL. Altogether, these results illuminate factors influencing donor-donor DNA rearrangement with bridge recombinases.

### Principles of improved human genome insertion specificity

To favor on-target insertion and mitigate DBL-only mediated recombination, we made three adjustments: (1) we encoded a unique bRNA recognition sequence not found in the genome (i.e. genome-orthogonal) on the delivered plasmid donor, (2) we used the enhanced version of the single bRNA (which limits donor-donor recombination) rather than the more active split bRNA system, and (3) we programmed the DBL to recognize the genome and the TBL to recognize the donor plasmid. To test this strategy, we selected 10 genomic loci that contain a unique 14 nt sequence close to their transcriptional start site and paired these sequences with a genome-orthogonal donor sequence, encoded within a puromycin-resistant plasmid (**Figure 3G**). We designed two enhanced single bRNAs for each sequence pair: one where the TBL recognizes the genome and the DBL recognizes the plasmid, and vice versa. We first evaluated the efficiency of the two orientations on day three post-transfection using ddPCR, observing a range of 0.18-3.38% for TBL-on-genome and 0.10-9.82% for DBL-on-genome (**Figure 3H**).

We next measured the specificity of these insertions after 18 days of puromycin selection. For 8 of 10 loci, we noticed that DBL targeting the genome was significantly more specific than TBL targeting the genome, reducing the number of off-targets by an average of 90.4% (range 87.0-94.1% reduction) (**Figure 3I, Figure S7A**). Despite confirming on-target insertion at each of the 10 loci three days post-transfection (**Figure 3H**), we observed on-target insertion for only four loci after 18 days of selection (**Figure 3J-M**), including for each of the top three most efficient loci (*SBF2*, *ANP32A*, and *IFNGR1*). We did not observe any correlation between insertion efficiency and specificity, indicating that the specificity gain with DBL is not an artifact of lower insertion efficiency (**Figure S7B**). Given the recovery of on-target insertions for the least efficient target (*DMD*), factors besides initial insertion efficiency such as differential fitness should be considered as well. Further analysis demonstrated that encoding a genome-orthogonal sequence on the plasmid donor additionally helped to eliminate plasmid-like off-targets for either bRNA orientation (**Figure 3N, Figure S7C-D**).

Overall, these results support our strategy for maximizing efficiency while balancing specificity (**Figure 3O**). We observed that recombination efficiency between a given pair of sequences can vary depending on which loop is programmed to which substrate, suggesting that both orientations should be tested for any gene of interest when optimizing for efficiency. Off-targets are consistently reduced by targeting the DBL instead of the TBL to the genome, an observation that is agnostic to the relative on-target insertion efficiency of the two loop orientations (**Figure S7H**). With DBL targeting of the genome, we achieved up to 82% on-target specificity (*SBF2*) and up to 9.8% efficiency (*ANP32A*). In conclusion, we propose that using a genome-orthogonal sequence on the donor plasmid, and programming the DBL to target the genome, is the optimal approach for performing genome insertion with IS622.

### Deep mutational scan of the IS622 recombinase

To further augment the effectiveness of IS622, we performed a deep mutational scan of the recombinase protein directly in human cells with the intention of bypassing variants that only improve activity in bacteria. To ensure compatibility with short read sequencing, we generated five libraries, each representing one-fifth of the recombinase, which included all possible mutations at each position (**Methods**). To incorporate these libraries into the genome, we first created a large serine recombinase (LSR) landing pad K562 cell line, which enables site-specific and single-copy integration, in contrast to random lentiviral integrations (Cao et al. 2021; Matreyek et al. 2017). This landing pad cell line was designed to capture IS622 recombinase variants at the AAVS1 safe harbor locus using our previously reported Dn29 LSR (Durrant et al. 2022), enabling high library coverage for robust screening (**Figure 4A, Figure S8A-G**). We transfected the resulting IS622 variant cell libraries with the enhanced bRNA and an inversion reporter, sorted them to isolate higher activity mutants, and then sequenced the recombinase in the pre- and post-sort populations (**Figure 4A**). We detected 99.6% of all possible single amino acid substitutions in the pre-sort cell population, and 51% of variants were enriched in the sorted population (**Figure S9A**). Randomized stretches of 65 aa as well as mutations of the recombinase catalytic residues were highly depleted in the sorted population, and less than 8% of mutations spread across 30% of residues were enriched above WT (**Figure 4B, Figure S9B**).

**Figure 4:**
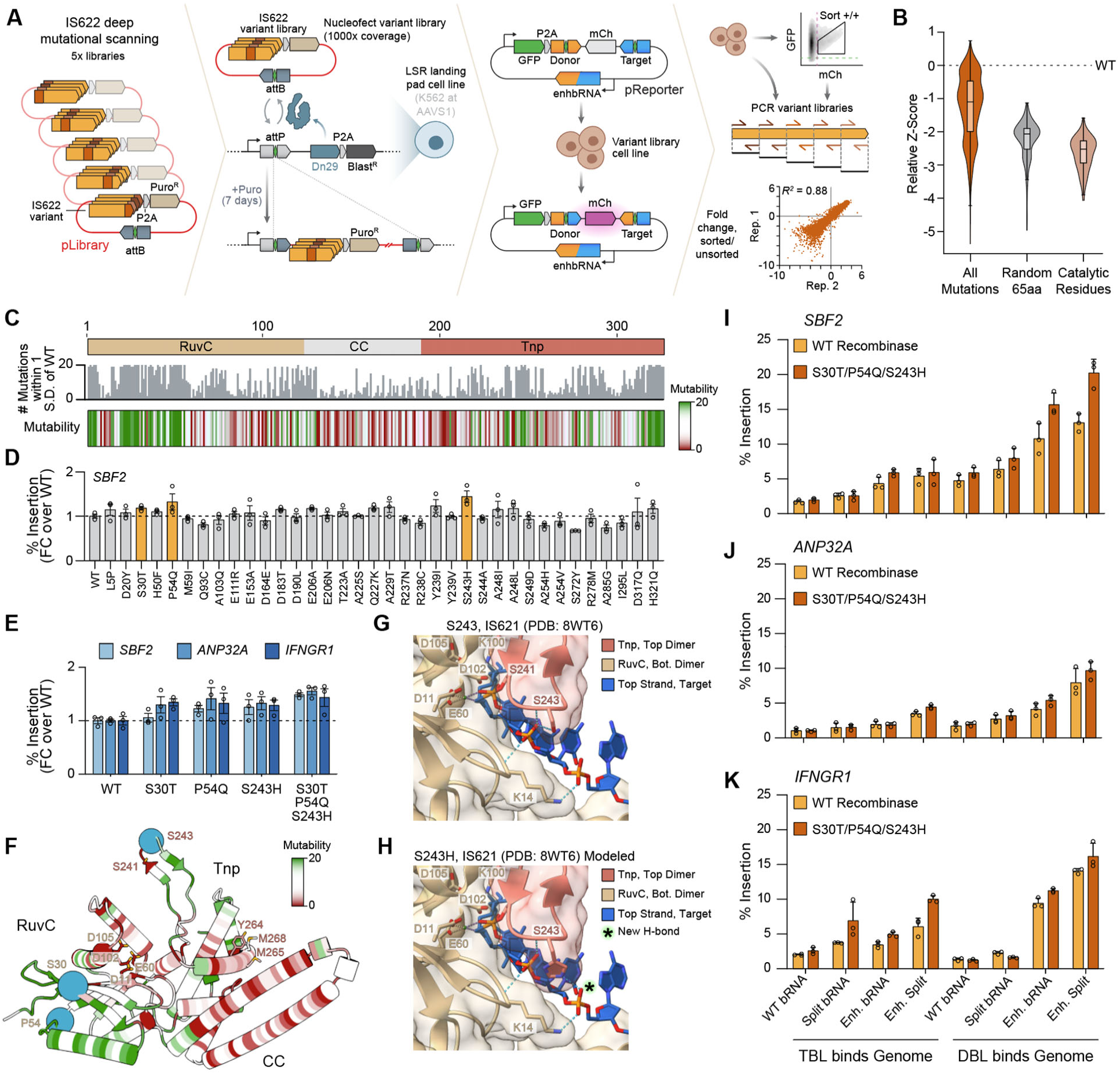
Improving IS622 activity by deep mutational scanning in human cells (**A**) Schematic of approach for a deep mutational scan of IS622 recombinase in human cells. Five libraries of single mutants are inserted into AAVS1 using Dn29 large serine recombinase. Genomically expressed recombinase uses a transfected enhanced bRNA to recombine an inversion reporter. Cells are sorted based on mCherry/GFP ratio prior to sequencing. (**B**) Z-score relative to WT for all mutations in deep mutational scan compared to random sequences serving as negative controls and to the catalytic residue positions (11, 60, 102, 105, and 241). (**C**) Mutability of the recombinase, represented as the number of mutations within one standard-deviation of the WT at each position. (**D**) Relative insertion efficiency at *SBF2* with selected single mutants. Insertion efficiency is measured using the enhanced bRNA with DBL binding the genome. (**E**) Relative insertion efficiency across genes with selected single mutants and a triple mutant. (**F**) AlphaFold 3 model of IS622 recombinase monomer. Color represents the per-position mutability score. Catalytic residues and wedge residues are highlighted as stick models, and residues mutated in the triple mutant are highlighted as sphere models. (**G**-**H**) DNA recognition by S243 in the IS621 structure (PDB: 8WT6) (**G**) and S243H in the IS621 model (**H**). Hydrogen bonds are depicted with blue dashed lines. (**I-K**) Genome insertion efficiency with WT and S30T/P54Q/S243H recombinase using various bRNA formats into *SBF2* (**I**), *ANP32A* (**J**), and *IFNGR1* (**K**) Enh., enhanced.

To interpret the mutational patterns, we developed a mutability score where we ranked each position of the recombinase according to how many amino acid substitutions per position fell within 1 standard deviation of the WT sequence (**Figure 4C, Figure S9C**). Based on this score, the RuvC domain is more mutable than the CC or Tnp domains, with the CC domain being least mutable overall. This finding is consistent with the critical roles of the CC domain in dimerization and the Tnp domain in bRNA binding (Hiraizumi et al. 2024). Based on the structural comparison between IS621 (PDB: 8WT6) and IS622 (AlphaFold3 model) (**Figure S3A-D**), we decided to use the IS621 structure to explore the likely solvent accessibility of specific amino acids in the context of the synaptic complex (**Methods**). We found that solvent-exposed residues were enriched in the screen (**Figure S9D**). Additionally, most mutations in the wedge residues of the Tnp domain were also depleted, consistent with their importance for DNA unwinding for base pairing with the bRNA (Hiraizumi et al. 2024). Based on both their mutability scores and fold-change activity, we selected variants and tested their activities for insertion into *SBF2* (**Figure 4D, Figure S9C**). Amongst these mutations, the S30T, P54Q, and S243H mutations consistently increased activity across genes (**Figure 4E**). We combined these mutations in a triple mutant, enhanced IS622, which exhibited consistent activity of 1.5-fold across genes (**Figure 4E**).

To interpret the potential mechanism of these activity enhancing mutations, we mapped the S30T, P54Q, and S243H mutations in IS622 onto the IS621 transpososome structure (PDB: 8WT6) (**Figure 4F**), as IS621 and IS622 share 88% sequence identity with no gaps and have the same number of residues (**Figure S2A**). P54 in IS622, equivalent to H54 in IS621, is likely located at a solvent-exposed loop, indicating that the P54Q mutation may improve solubility or aid in protein folding. In the IS621 structure, T30 hydrogen bonds with U142 of the DBL, stabilizing the tetrameric complex (Hiraizumi et al. 2024) (**Figure S10A**). It seems likely that S30 of IS622 forms a similar hydrogen bond with the DBL, and that the S30T mutation enhances activity due to a favorable additional contact between the methyl group of T30 and the adjacent D20 residue (**Figure S10B-C**). The catalytic site of IS621 is created only upon tetramerization, and consists of the RuvC domain from one dimer and the Tnp domain from the opposing dimer (Hiraizumi et al. 2024). The catalytic S241 forms a covalent bond with DNA, while the main chains of K14 and S243 interact with the DNA phosphate backbone at the catalytic site (**Figure 4G**). Modeling of S243H suggests that the H243 side chain forms a new hydrogen bond with the DNA phosphate backbone (**Figure 4H**), thereby increasing activity.

In the validations of our deep mutational scan, we evaluated mutations using the enhanced bRNA, where the DBL was programmed to bind the genome. To assess if the enhanced IS622 recombinase was generalizable to other bRNA formats, we evaluated either bRNA orientation with various bRNA scaffolds (**Figure 4I-K**). Across all conditions, increased activity was consistently observed with the enhanced bRNA scaffolds in comparison to the unmodified bridge RNAs. Pairing these with the engineered recombinase yielded up to 20.2% insertion at *SBF2*. Across three targets, we show that the original bridge recombination system (using WT bRNA where the TBL binds the genome) can be outperformed 9.8-fold by utilizing the optimal loop configuration, our enhanced bRNA scaffolds, and our engineered recombinase.

### Megabase-scale genome rearrangements with bridge recombination

Genomic insertion has been achieved with many natural and engineered molecular systems (Tou & Kleinstiver 2022; Villiger et al. 2024). Some of these tools have been co-opted for genomic rearrangement, but none of them are naturally capable of this function. Bridge recombinases are unique in that they can be programmed to recognize almost any pair of genomic sequences, without the pre-installation of recombinase recognition sites common among other approaches (Anzalone et al. 2021; Yarnall et al. 2022).

To demonstrate this capability in human cells, we took a similar approach to our genome insertion assays. We first nominated pairs of unique sequences in the genome, filtering for matching core sequences and non-matching positions 6 and 7 (**Methods**). Depending on the relative strands of these two sequences, recombination between the two sites yields intra-genomic inversion or excision, which can be measured by tagmentation and amplicon sequencing (**Figure 5A**). The effect of loop orientation on insertion efficiency prompted us to assess both configurations of the enhanced bRNA for these rearrangements (site 1 bound by TBL, site 2 bound by DBL, and vice-versa). We measured recombination efficiency for seven pairs of sequences resulting in inversion (**Figure 5B**), and seven pairs of sequences resulting in excision (**Figure 5C**). We observed up to 10.3% efficiency of inversion (1,153 bp) and 12.8% efficiency of excision (55,840 bp). The efficiency of recombination had no clear correlation with the distance between the recombined sites, and we were able to invert as long as 929,524 bp (0.92 Mbp, 2.56% efficiency) and excise as long as 134,143 bp (0.13 Mbp, 1.10% efficiency).

**Figure 5:**
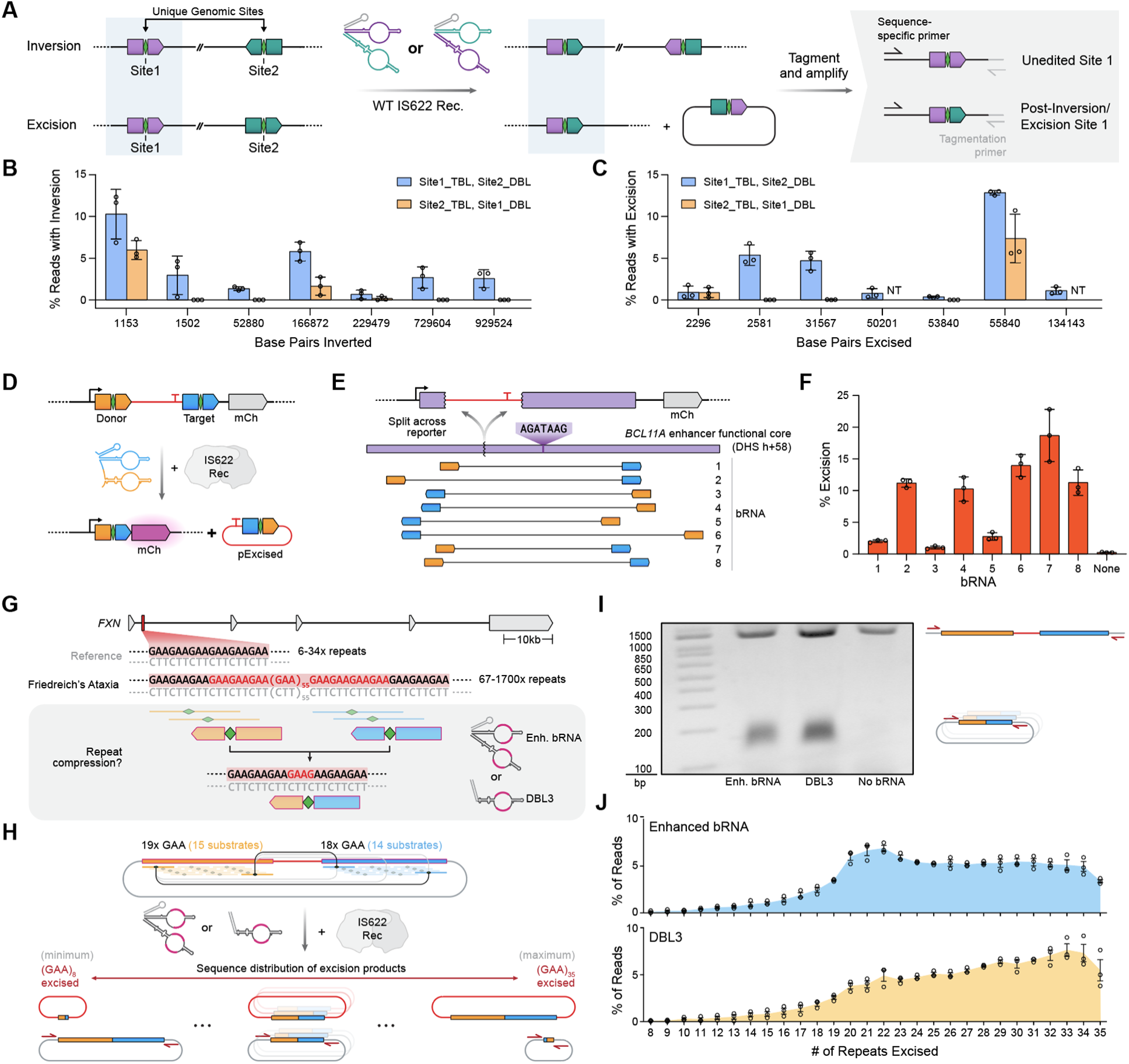
Applications of programmable genomic rearrangement with IS622 (**A**) Schematic of performing intra-chromosomal recombination. bRNAs of both orientations are used to yield inversion or excision, which is measured by tagmentation and next-generation sequencing. (**B**) Efficiency of genome inversion across 7 sequence pairs. Each pair of bars represents one pair of sequences recombined with either orientation of bRNA. (**C**) Efficiency of genome excision across 7 sequence pairs. Each pair of bars represents one pair of sequences recombined with either orientation of bRNA. NT, not tested. (**D**) Schematic of excision reporter assay. Removal of a terminator ahead of the promoter yields mCherry expression. (**E**) Schematic of bRNAs used to recombine the *BCL11A* enhancer region on the plasmid excision reporter. (**F**) Efficiency of *BCL11A* enhancer excision with eight unique bRNAs. (**G**) Overview of *FXN* repeat expansion and compression via bridge recombination. Enh., enhanced. (**H**) Schematic of plasmid reporter for recombination of repeat regions. Possible recognition sites are represented by colored lines, with each green diamond representing the core of a sequence. Recombination results in a distribution of product lengths. (**I**) Agarose gel showing efficiency of repeat recombination with 1) an enhanced (Enh.) bRNA and 2) a DBL (DBL3) in comparison to 3) no bRNA. Lower band represents the range of remaining repeats. (**J**) Distribution in the number of repeats excised, measured via next-generation sequencing.

Across both rearrangement types, we observed that the bRNA orientation again influences the outcome. For 7 of the 12 sequence pairs where we tested both orientations, only one orientation mediated recombination, with the other yielding little to no rearrangement (0-0.03%). For the remaining 5 sequence pairs, we observed recombination with both orientations at similar efficiency (relative fold-change of 1.0-3.8). These results suggest that the main driver of rearrangement efficiency is intrinsic to the recombinase and bRNA, rather than to the specific pair of sequences and their respective genomic contexts.

### BCL11A enhancer excision with bridge recombination

The ability to modify the genome in a sequence-specific manner is a proven method for interrogating genotype-phenotype relationships (Adamson et al. 2016; Dixit et al. 2016; Shalem et al. 2015; Wang et al. 2014). In particular, CRISPR functional genomics has unveiled a number of new disease targets (Przybyla & Gilbert 2022), some of which have led to the development of effective therapies. Prominently, an enhancer (DHS h+58) of the transcription factor gene *BCL11A* was found to be essential to its expression in a CRISPR screen (Canver et al. 2015), and targeted knockout of this region in erythroid cells is now used as a therapy for sickle cell anemia and beta-thalassemia (Frangoul et al. 2020).

The ability of bridge recombination to programmably excise specific genomic intervals represents a significant opportunity for high-resolution functional genomics. As a proof-of-concept, we synthesized the *BCL11A* enhancer region with a disrupting intervening sequence that must be excised from a reporter plasmid in order to express mCherry, allowing us to measure excision efficiency (**Figure 5D-E**). This region encodes a GATA motif, elimination of which attenuates *BCL11A* expression in erythroid progenitors (Canver et al. 2015). Across a variety of binding sites for both the TBL and DBL, we were able to achieve robust excision of up to 18.7%, indicating that bridge excision can be applied at user-defined intervals (**Figure 5F**).

### Excision of Friedreich’s ataxia repeat expansion

Repeat expansion disorders are a class of ∼50 genetic diseases arising from the accumulation of repetitive sequences in the genome. These repeats are typically within coding genes and cause disease via loss-of-function, dysregulation of gene expression, generation of toxic RNAs, or expression of gain-of-function proteins (Khristich & Mirkin 2020; Malik et al. 2021). Among these diseases is Friedreich’s ataxia, a neuromuscular disorder resulting from the expansion of a GAA trinucleotide repeat in intron 1 of the gene *FXN* (**Figure 5G**). In healthy individuals, there are as few as 6 repeats, while patients exhibiting the disease encode as many as 1700 repeats (Bürk 2017).

Using bridge recombinases to excise excess repeats should be scarless and in-frame, enabling the restoration of a corrected genotype. However, this type of recombination would require both the TBL and DBL to bind identical substrates, potentially causing competitive inhibition of one loop by the other. To this end, we wondered if DBL-only recombination (**Figure 3F**) could be exploited for the removal of undesired repeat regions. To test this, we programmed both loops of enhanced bRNA or an enhanced DBL (DBL3) to recognize a 14 nt sequence spread across 5 GAA repeats. Within just 40 GAA repeats, there are 36 stretches of this 14 nt recognition sequence, with each CT core on the bottom strand representing a unique recombination breakpoint. Recombination can occur between any two of these potential binding sites, all of which result in the excision of repeats, leading to variable outcomes across different molecules (**Figure 5G**).

We encoded two GAA repeat regions within a plasmid on either side of a stuffer sequence to compare excision efficiency between a full bRNA and a DBL alone (**Figure 5H**). We amplified across this region in order to quantify the proportion of intact and excised reporters (**Figure 5I**). Since the total number of repeats is related to the severity of disease (Khristich & Mirkin 2020), we sought to quantify how many repeats were removed from the plasmid substrates. As expected, a range of outcomes was observed for both bRNA and DBL-only, with as few as 8 and as many as 35 repeats excised from the plasmids (**Figure 5J**). Notably, the DBL appeared to be more effective at excising a larger number of repeats, with 40% of plasmids exhibiting excision having 80% (>29 repeats) or more of the repeats removed. These results show that genomic excision with bridge recombination simply requires the 326 aa recombinase (976 nt) and 90 nt DBL to mediate excision. Future translational efforts may be enabled by the transient delivery of these components as RNA molecules, facilitating genomic modification without any DNA delivery.

## DISCUSSION

In this study, we present IS622, the first bridge recombinase for programmable insertion, inversion, and excision of the human genome. We interrogate the mechanism and application of IS622 through systematic engineering, enhancing activity of both its bridge RNA and recombinase. We highlight key strategies for maximizing the effectiveness of applying bridge recombination in human cells, particularly the use of the donor-binding loop to base-pair with the genome, revealing functional and mechanistic differences between the two bRNA binding loops. Our optimized approach for universal chromosomal DNA rearrangements achieves as high as 20% insertion efficiency and mobilizes genomic DNA at the megabase scale.

As the applications for DNA recombinases have grown increasingly ambitious over the past few decades, the need to overcome the limitations of existing tools has also grown. Recombinase platforms have advanced beyond early implementations of Cre-LoxP knockouts (Orban et al. 1992; Soriano 1999) to sophisticated applications in studying development (Chow et al. 2021), building biological circuitry (Merrick et al. 2018; Weinberg et al. 2017), and developing gene therapies (Pandey et al. 2024; Xie et al. 2024). These approaches have been developed through many systematic engineering and discovery efforts to tackle challenges such as recognizing user-defined sequences, minimizing scars, or targeting to safe-harbor loci (Durrant et al. 2022; Fanton et al. 2024; Fauser et al. 2024; Jelicic et al. 2023; Schmitt et al. 2016). The bispecific and programmable mechanism of bridge recombinases naturally circumvents many of the fundamental engineering challenges in the recombinase field. We expect future efforts for improving bridge recombinases to focus on further improvement of targeting efficiency and specificity, delivery of megabase-scale DNA payloads, and development of therapeutic delivery formulations, including RNA-only delivery for genomic rearrangements such as repeat excisions.

Bridge recombinases can modify the genome at arbitrary new scales, ranging from single gene insertions to megabase-sized rearrangements, which unlocks significant potential for understanding cellular function and human disease pathology. Locus- and chromosome-scale excisions or inversions will enable higher resolution understanding of sequence-to-function relationships across organisms. In combination with chromosomal translocation, these recombinations can enable precise creation of cell lines and animal models which mimic the genotypes of cancers and other chromosomal abnormalities. Finally, bridge recombination is poised to be combined with AI-generated DNA sequences of high complexity (Brixi et al. 2025; Nguyen et al. 2024), enabling programmable genome design at unprecedented length scales.

## SUPPLEMENTARY FIGURES

**Figure S1:**
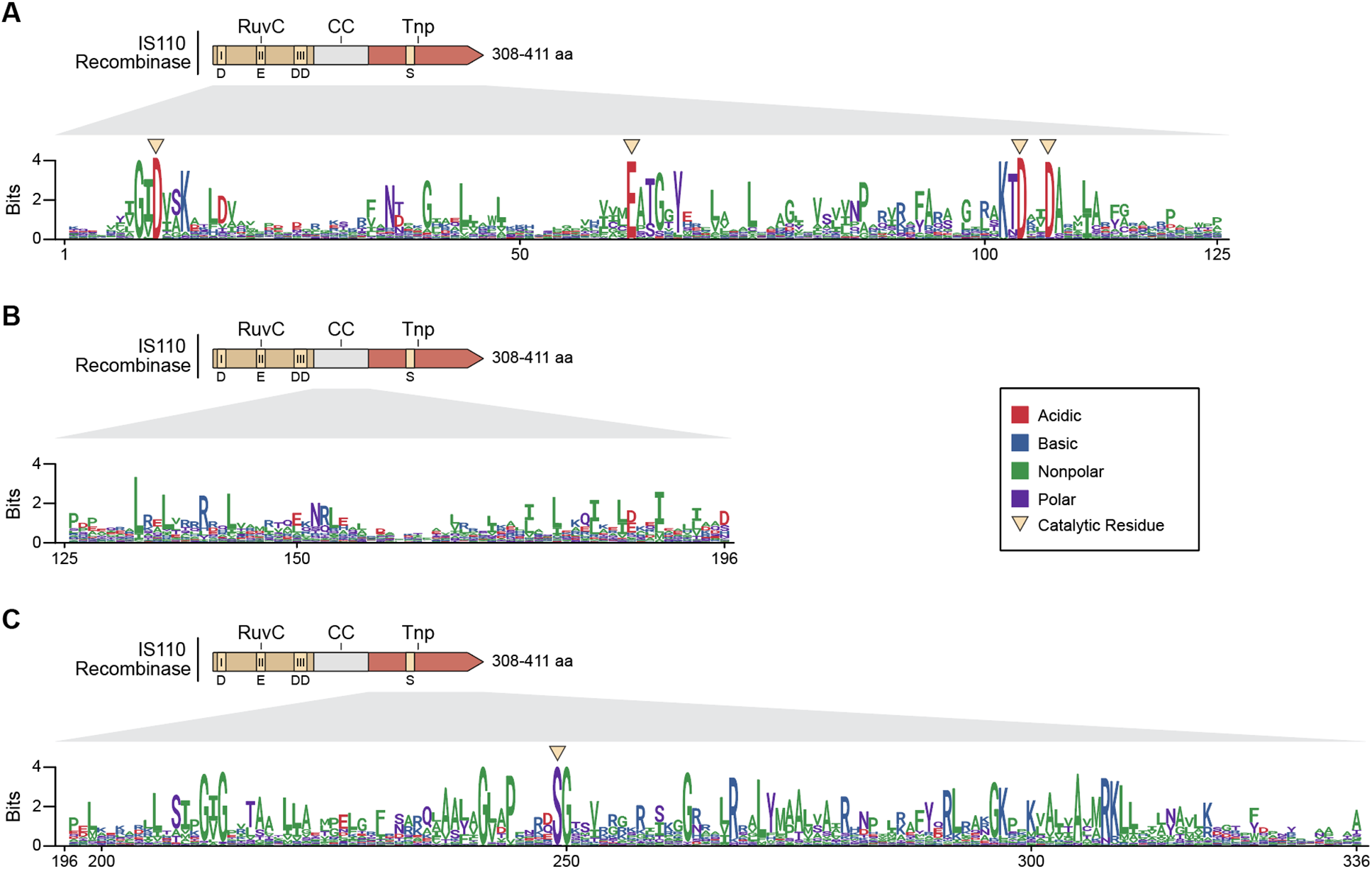
Diversity of tested bridge recombinases (**A-C**) Sequence conservation of 72 bridge recombinases in the RuvC domain (**A**), CC domain (**B**), and Tnp domain (**C**). Alignment columns with more than 75% gaps are filtered out for clarity.

**Figure S2:**
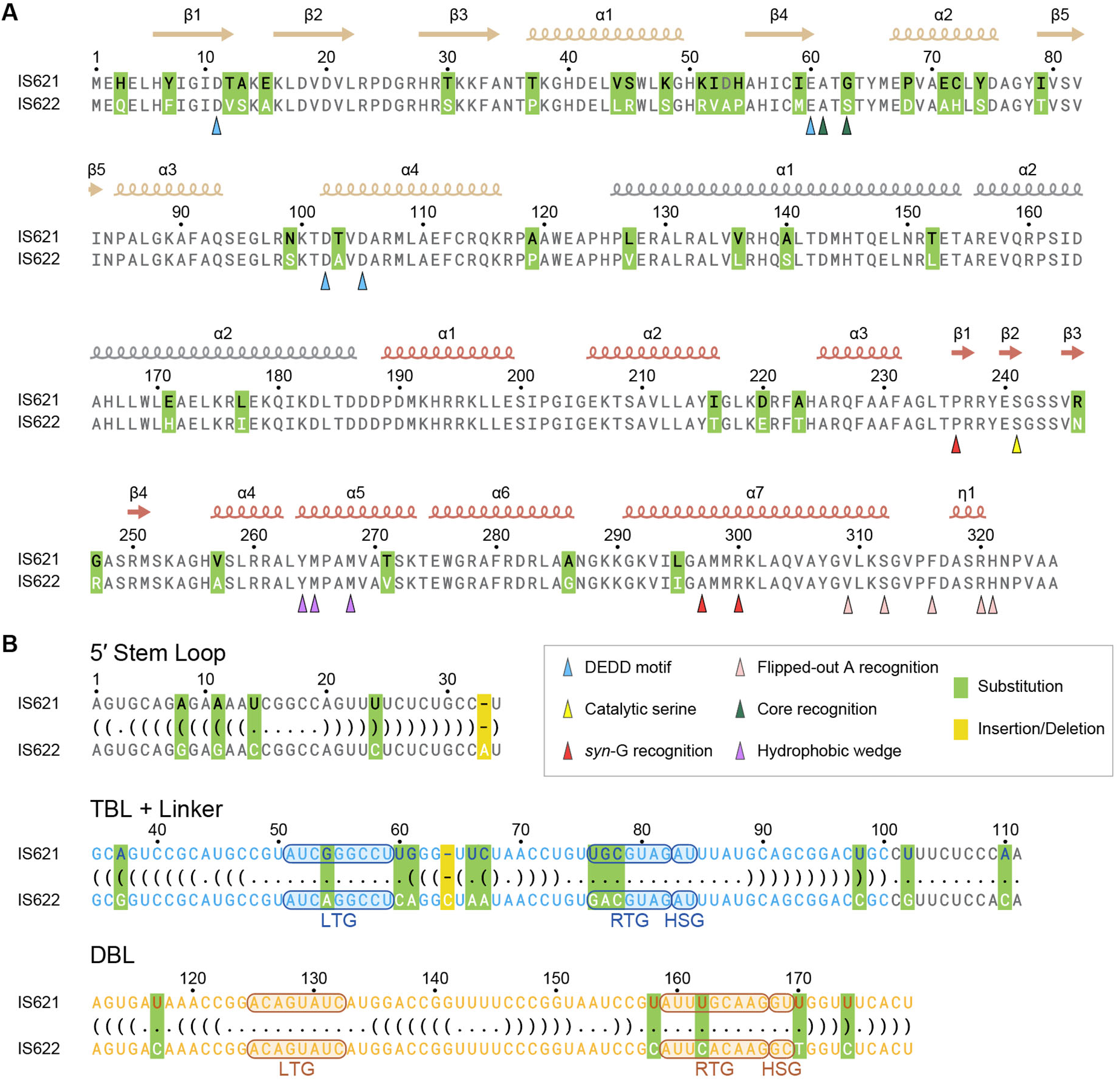
Sequence comparison between IS621 and IS622 (A) Alignment of IS621 and IS622 recombinases. Secondary structures are shown above the sequences for IS621 (PDB: 8WT6). Differences are highlighted in green. (**B**) Alignment of IS621 and IS622 bRNAs. Dot-bracket notation of bRNA structure based on IS621 bRNA structure. Differences highlighted in green and gold.

**Figure S3:**
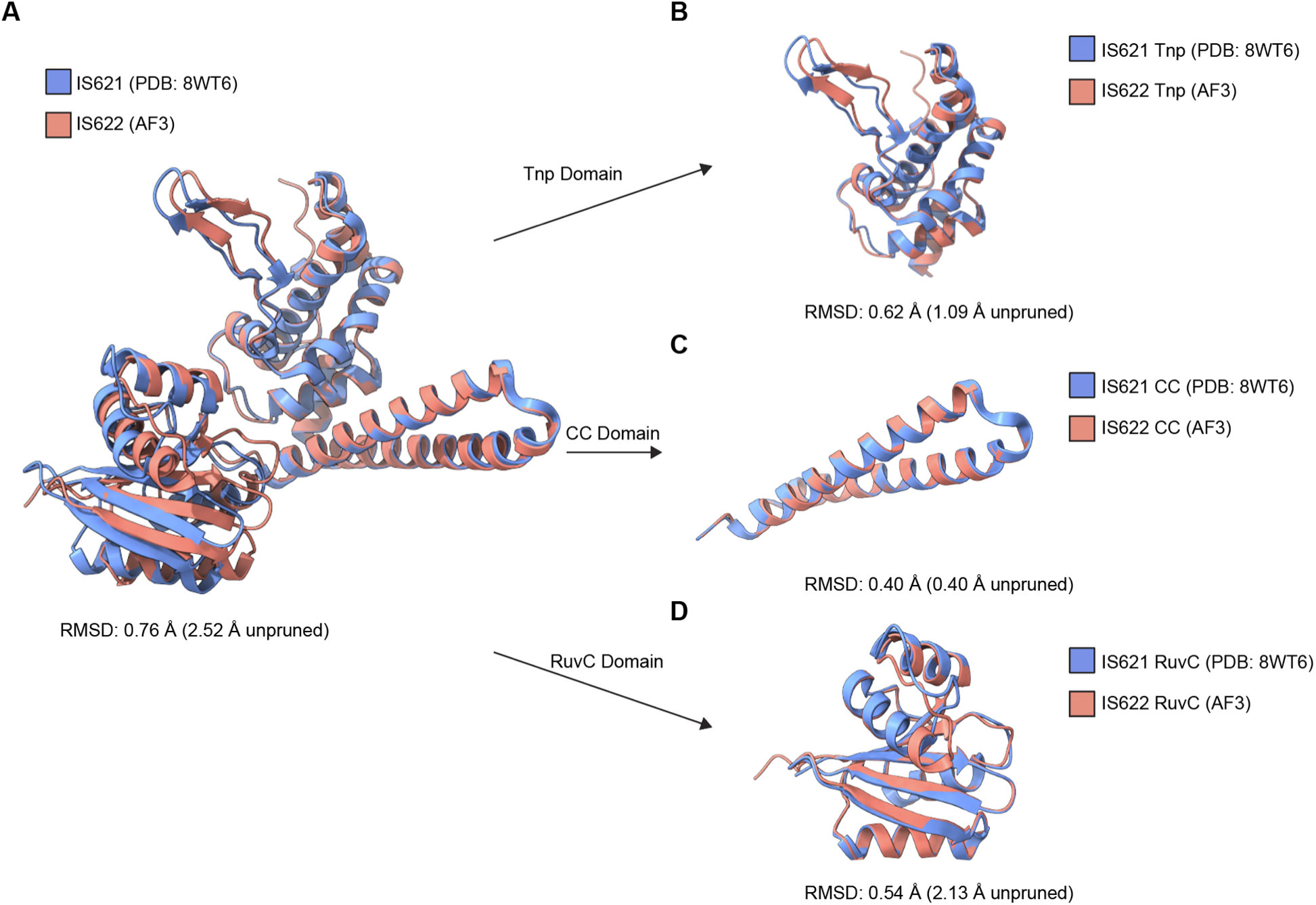
Comparison of the IS621 structure with the predicted model of IS622 (**A**) Structural alignment of IS621 recombinase monomer (PDB: 8WT6) with IS622 recombinase monomer predicted with AlphaFold 3. (**B-D**) Structural alignment of individual domains of IS621 and IS622 for the Tnp domain (**B**), CC domain (**C**), and RuvC domain (**D**).

**Figure S4:**
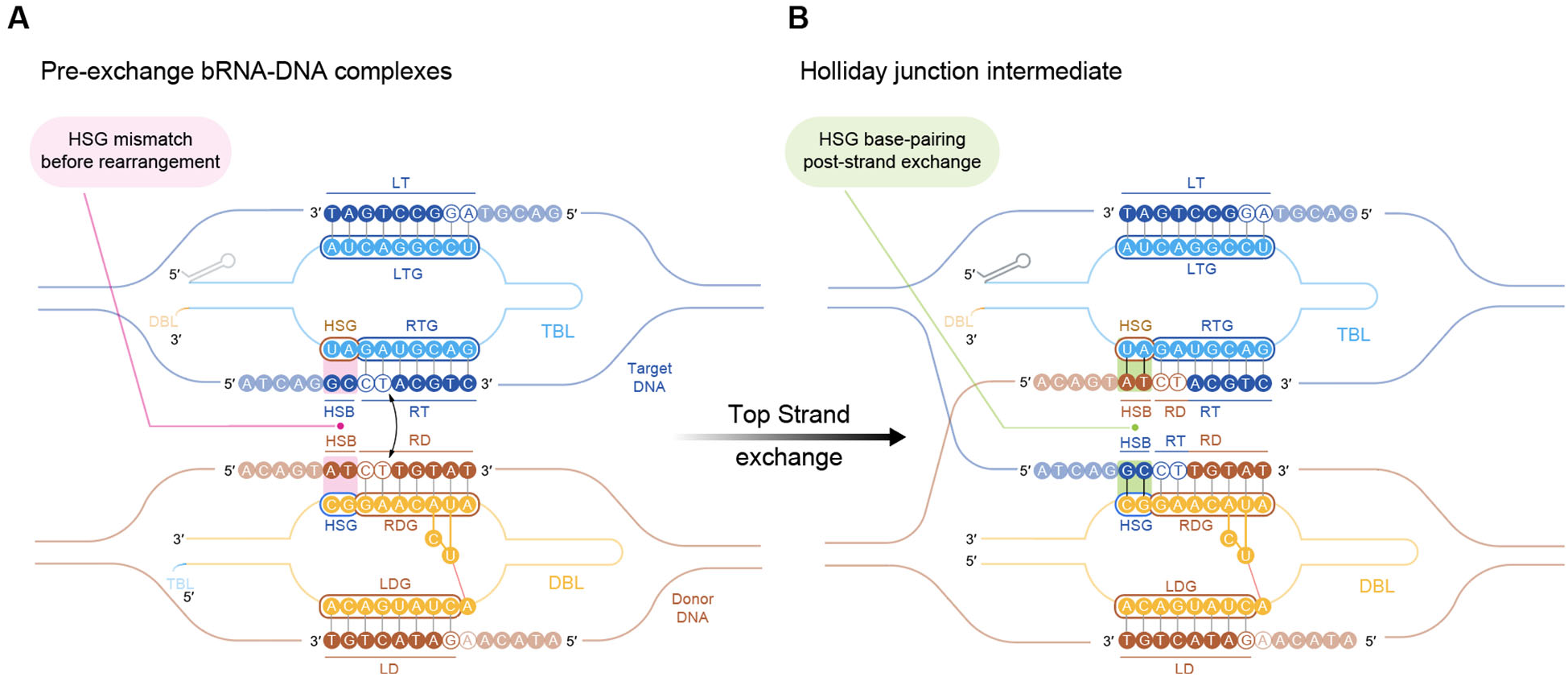
Handshake guide base-pairing mechanism (**A-B**) Model of TBL and DBL base-pairing with target and donor, pre- (**A**) and post- (**B**) strand exchange. Handshake guides mismatch with DNA upstream of core sequences prior to strand exchange and base pair with DNA after strand exchange. LTG, left target guide; RTG, right target guide; LDG, left donor guide; RDG, right donor guide; TBL, target-binding loop; DBL, donor-binding loop; HSG, handshake guide; HSB, handshake bases.

**Figure S5:**
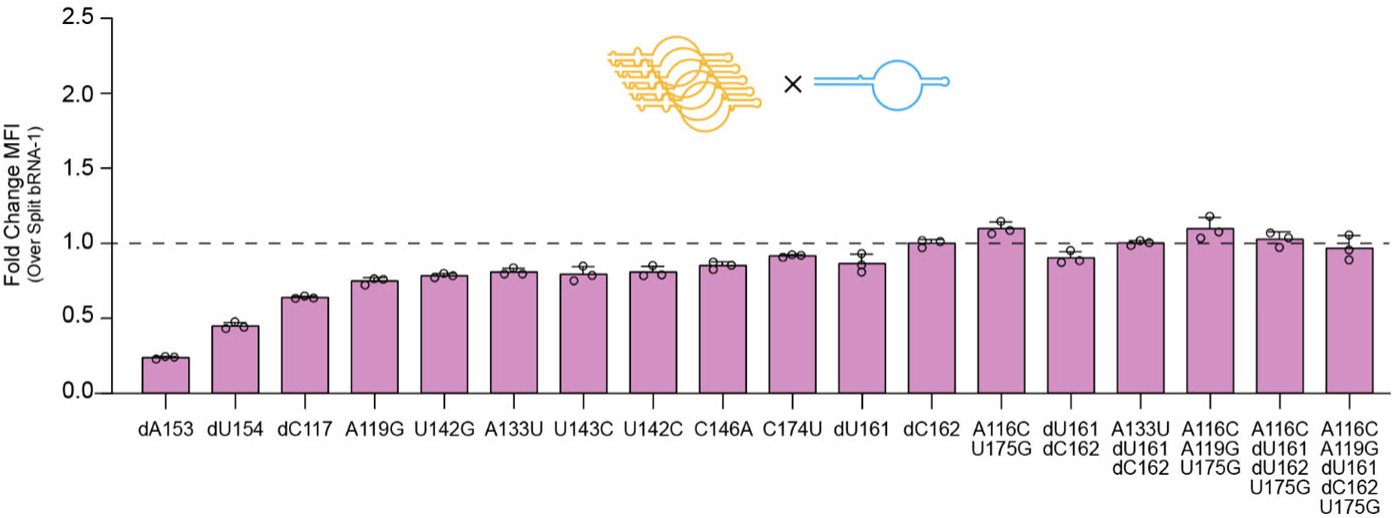
Donor binding loop rational mutagenesis Efficiency of recombination of DBL mutants in inversion reporter assay. DBL mutants tested with boundaries of 112-179 and paired with a TBL with boundaries of 35-100.

**Figure S6:**
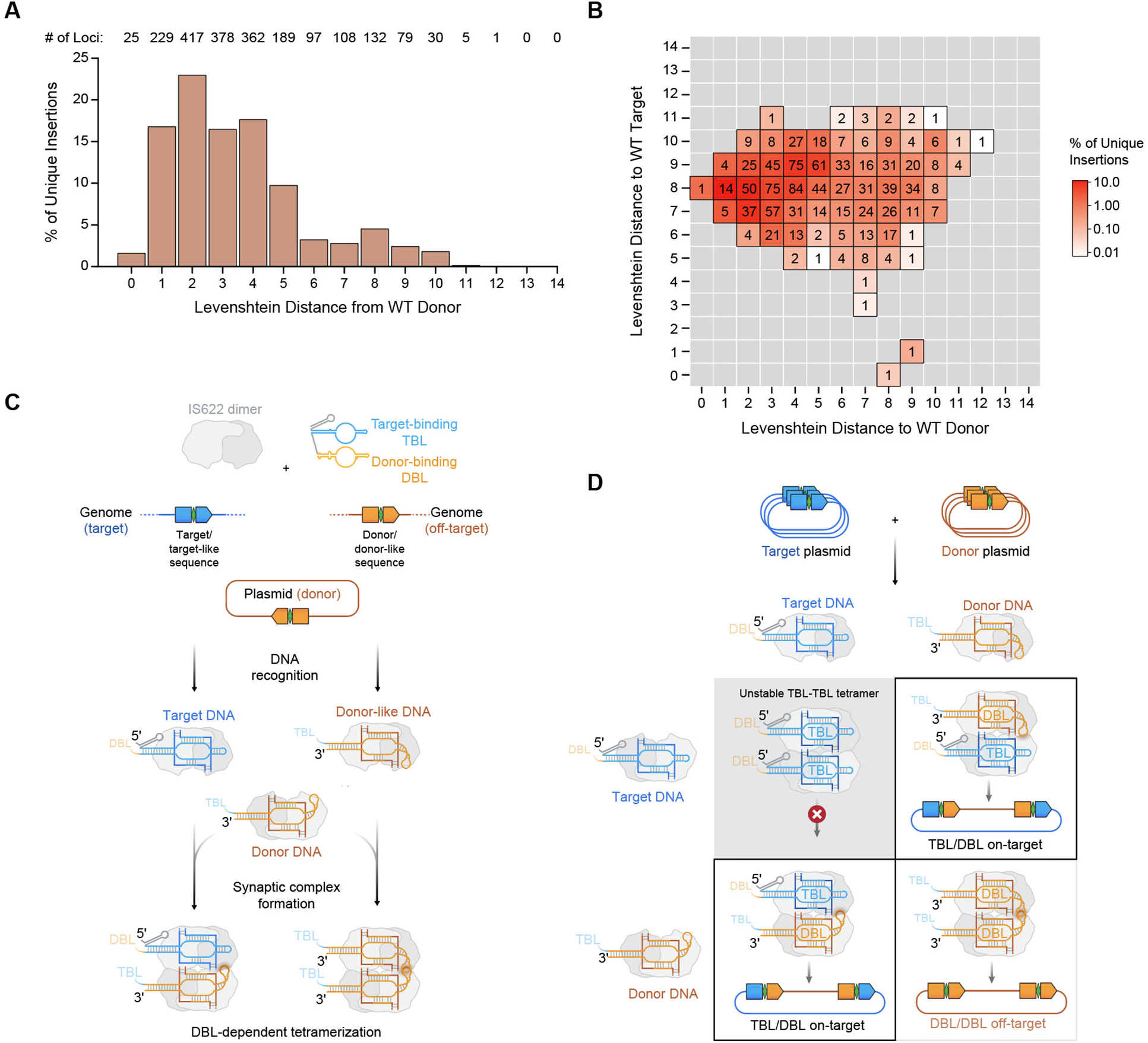
Assessment and mechanism of DBL-DBL guided off-targets (**A**) Levenshtein distance of off-targets from the use of a WT IS622 bRNA with a WT donor plasmid. Percentage of unique insertions based on the total number of recovered UMIs across all insertions genome wide. The number of loci represented within each bar is shown at the top. (**B**) Heatmap of Levenshtein distance of off-targets in comparison to WT target and WT donor. The number of loci represented are shown in the center of each cell. (**C**) Mechanism of tetramerization with one TBL and one DBL or between two DBLs in the context of human genome insertion. (**D**) General mechanism of possible dimer-dimer interactions between DNA-bound TBL and DBL dimers. TBL-TBL tetramers are destabilized by the lack of a long hairpin reaching between dimers.

**Figure S7:**
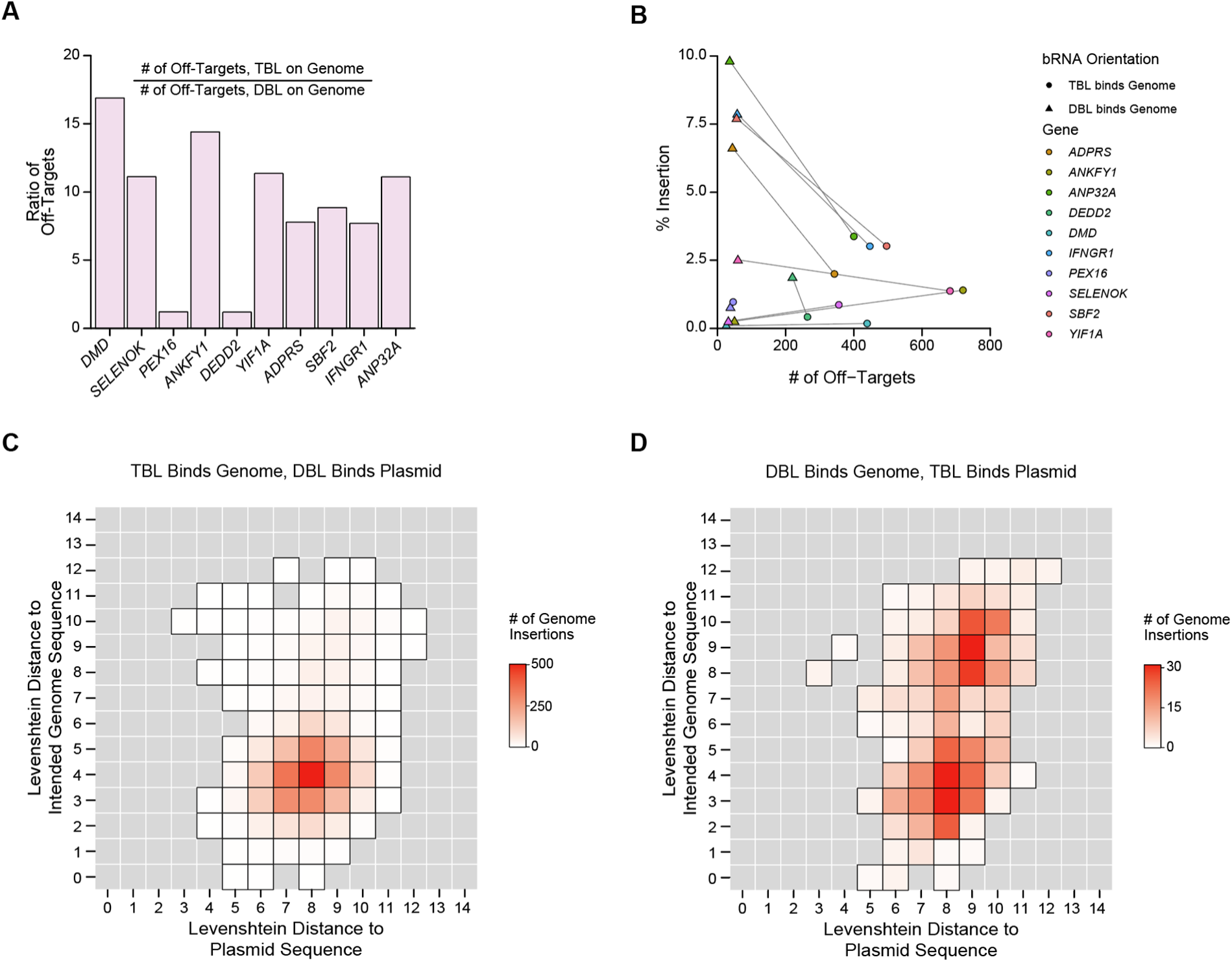
Specificity and off-target analysis of programmable gene insertion (**A**) Ratio of the number of off-targets when the bRNA is designed for the TBL to bind the genome versus for the DBL to bind the genome. (**B**) Comparison of efficiency and number of off-targets across all genes between TBL or DBL bound to the genome. (**C**) Heatmap of Levenshtein distance of off-targets in comparison to expected genome site or plasmid encoded sequence when the TBL is encoded to bind the genome. (**D**) Heatmap of Levenshtein distance of off-targets in comparison to expected genome site or plasmid encoded sequence when the DBL is encoded to bind the genome.

**Figure S8:**
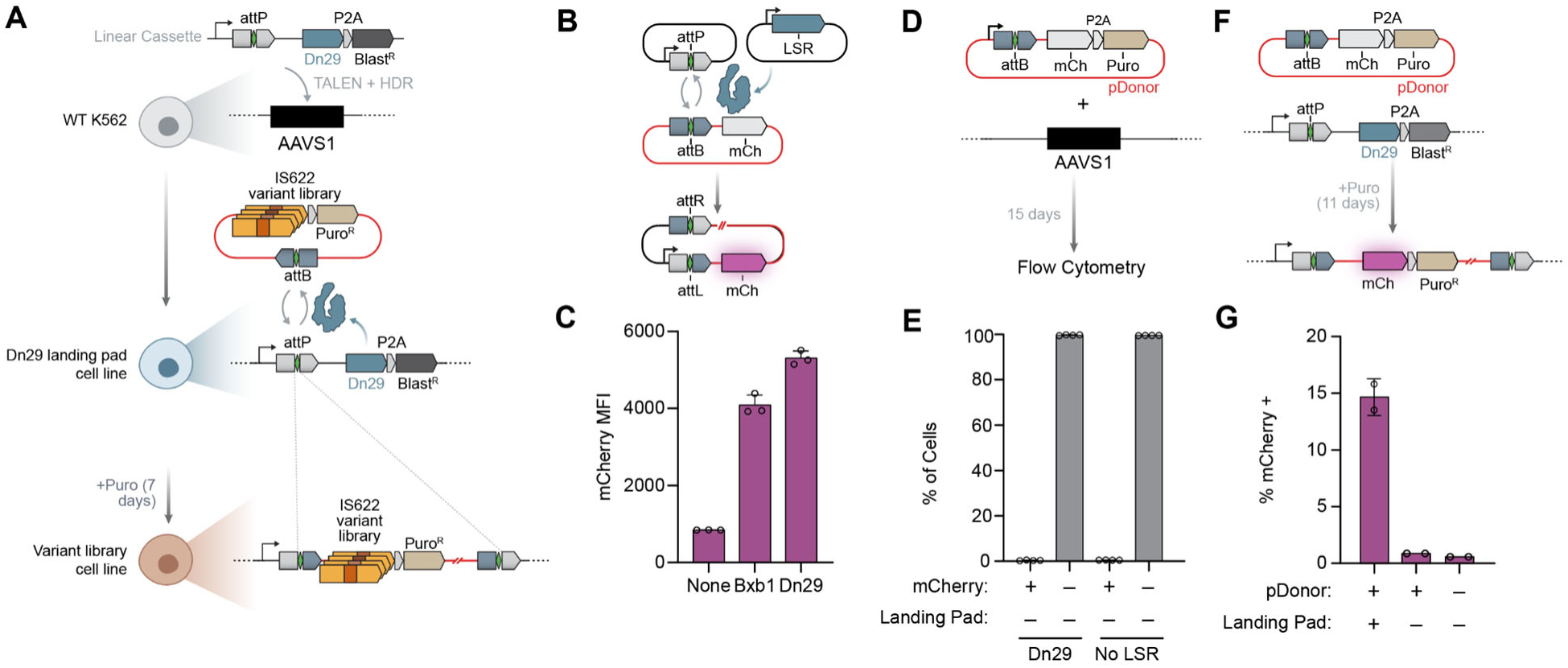
Generation and validation of Dn29 landing pad cell line (**A**) Schematic of cell line creation. attP-Dn29-P2A-Blast^R^ was knocked into AAVS1 using TALENs. AttB-containing plasmids are subsequently knocked into the AAVS1 locus via Dn29 mediated attP/attB recombination. (**B**) Schematic of plasmid recombination assay for evaluating LSR efficiency. (**C**) Relative recombination efficiency of Bxb1 and Dn29 LSRs. (**D**) Schematic of off-target insertion assay. AttB-containing plasmids were nucleofected into WT K562 cells, and mCherry expression was measured after 15 days in culture. (**E**) Percentage of WT K562 cells bearing mCherry knock-in after 15 days. (**F**) Schematic of landing pad cell line insertion efficiency assay. mCherry-P2A-Puro^R^ is knocked into the genome and cultured under puromycin selection for 11 days. (**G**) Percentage of Dn29 landing pad cells with mCherry knock-in after 15 days.

**Figure S9:**
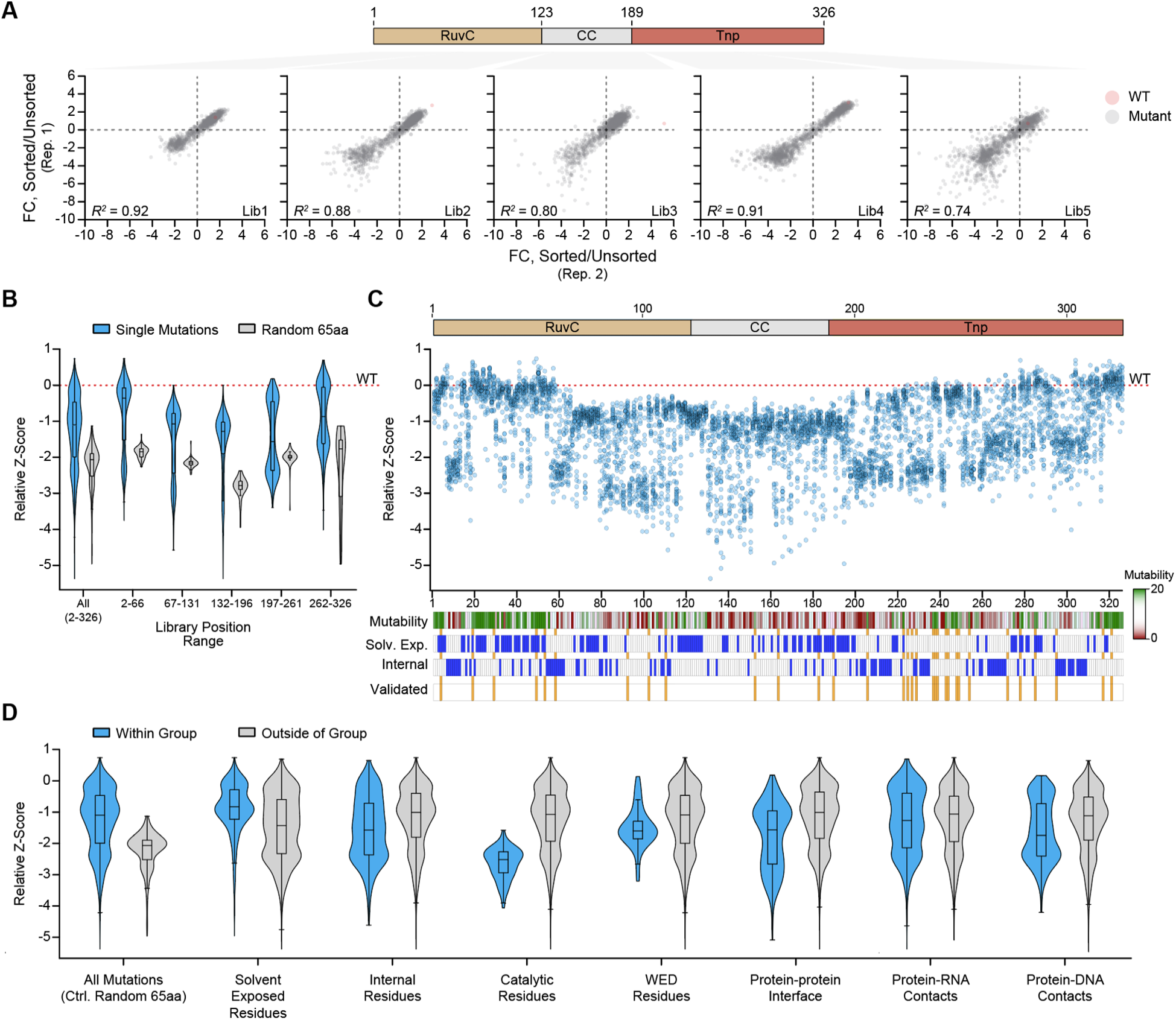
Deep mutational scan distributions and mutant validations (**A**) Replicate correlation for each 65 aa segment of deep mutation scan. WT recombinase highlighted in red. FC, fold-change. (**B**) Distribution of all mutations compared to distributions of random 65 aa negative controls. (**C**) Distribution of all mutations per position. Mutability score, solvent exposure, internal, and validated residues are highlighted. (**D**) Distribution of mutations within and outside of various residue groups. Groups determined based on IS621 transposome structure (PDB: 8WT6).

**Figure S10:**
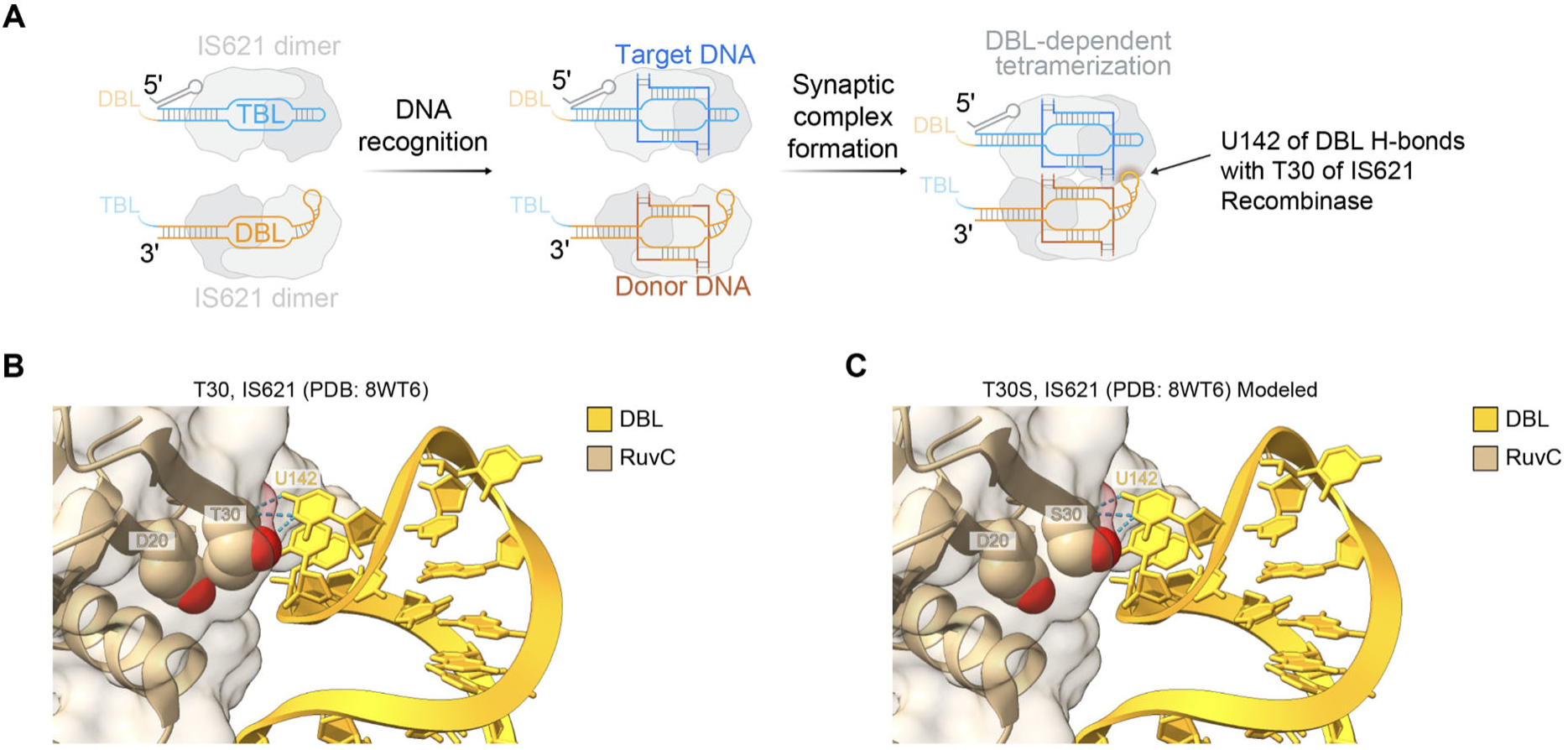
Modeling of the T30S mutation on the IS621 structure (**A**) Overview of bridge recombinase synaptic complex assembly, based on IS621. TBL, target-binding loop; DBL, donor-binding loop. U142 of the DBL contacts S30 of the IS621 recombinase. (**B-C**) Recognition of the DBL U142 by T30 (**B**) and T30S (model) (**C**) in the IS621 structure (PDB: 8WT6). Contacts between D20 and X30 (T30 or S30) are compared.

## METHODS

### Cell lines and culture

Experiments were conducted in HEK293FT and K562 cells. HEK293FT cells were cultured at 37°C with 5% CO_2_ in DMEM + GlutaMax supplemented with 10% FBS and 100 U/mL PenStrep. K562 cells were cultured at 37°C with 5% CO_2_ in RPMI + GlutaMax supplemented with 10% FBS and 100 U/mL PenStrep.

### Bridge recombinase ortholog mining

Bridge recombinase candidates were identified from our previously generated database of assembled genomic and metagenomic sequences, focusing on those that were within the broader IS621-like clade (Durrant et al. 2024). We identified bRNA sequences in IS110 loci using cmsearch (INFERNAL v1.1.4) and an RNA covariance model that was constructed using the IS621 bRNA as a seed sequence. The search was performed using the --toponly parameter as all loci were oriented w.r.t. the forward strand of the bridge recombinase coding sequence. Bridge RNA sequences were filtered by an E-value threshold of 1e-6 and by requiring at least 80% coverage of the covariance model length.

### Ortholog screening in human cells

The recombinase, bRNA, target, and donor sequences for each nominated ortholog were cloned into plasmids for evaluating activity. For orthologs with >75% amino acid identity to IS621, positions 12-14 of the target were mutated to match the target binding loop RTG. Bridge recombinases were human codon optimized and then cloned into the same construct as the bRNA. The bRNA is expressed via a U6 promoter, and the recombinase is expressed via an Ef1a promoter. The recombinases were cloned with a C-terminal NLS as Rec-3xSV40NLS-P2A-EGFP. The target and donor were cloned on a plasmid separated by 1000 bp. 100 ng of recombinase and bRNA plasmid was cotransfected with 288 ng of target + donor plasmid into 1.4 x 10^4^ HEK293FT cells using 0.5 µL Lipofectamine 2000 in 96-well plates. After 60 hours at 37°C, plasmid DNA was recovered from cells using 50 µL of QuickExtract (Biosearch Technologies) per well. Extracted DNA was cleaned up using 0.9x AmpureXP magnetic beads (Beckman Coulter). PCR was performed with 2x Platinum SuperFi II across the newly formed inversion junction. Activity was determined via agarose gel electrophoresis of the PCR and subsequent visual inspection.

### Luciferase reporter assay in human cells

100 ng of recombinase and bRNA plasmid was cotransfected with 288 ng of C-luciferase/G-luciferase inversion reporter plasmid into 1.4 x 10^4^ HEK293FT cells using 0.5 µL Lipofectamine 2000 in 96-well plates. C-luciferase was expressed constitutively from the CMV promoter. G-luciferase was flanked by the bRNAs natural target and donor sequence, or, a target and donor sequence cognate for the bRNA binding loops. Inversion of G-luciferase results in in-frame expression of the luciferase from an Ef1a promoter. After 60 hours at 37°C, 20 µL of cell media from each well was transferred to two different 96-well black flat bottom plates. Luciferase expression was measured using the luciferase activity kit from Targeting Systems. G-luciferase expression was measured by adding 50 µL of GAR diluted 1:100 in GAR buffer to cell media, incubating in the dark for 10 minutes, then measuring with luminescence on Tecan with 10 ms exposure time. C-luciferase expression was measured by adding 50 µL of VLAR diluted 1:100 in VLAR buffer to cell media, incubating in the dark for 10 minutes, then measuring with luminescence on Tecan with 10ms exposure time. Inversion activity was measured as the ratio of G-luciferase luminescence divided by C-luciferase luminescence, yielding a measurement in relative fluorescence units (RLU). Inversion was confirmed via amplification of the inversion junction on the luciferase reporter plasmid. For 161642, IS621, 127209, and IS622, the natural target sequence was modified such that positions 12-14 matched the target binding loop RTG.

### AlphaFold modeling of IS622 recombinase and alignments

The three-dimensional structure of IS622 recombinase was predicted using the AlphaFold 3 (Abramson et al. 2024) Server from the amino acid sequence alone and modeled as a single monomer. From the five models outputted, the resulting model was selected for analysis based on the minimal RMSD to one of two monomers with resolved catalytic loops from the experimentally determined structure of IS621 (PDB ID: 8WT6) using UCSF ChimeraX 1.9 (Pettersen et al. 2021). Alignments were performed using the matchmaker function with backbone atoms for the three functional domains alongside the whole protein.

### IS622 bRNA engineering

The activity of IS622 engineered bRNAs was measured in HEK293FT cells. In the case of a single bRNA, 100 ng of a plasmid encoding U6-bRNA and Ef1a-Rec-3xSV40NLS-P2A-EGFP was cotransfected with 288 ng of a plasmid encoding Efa-Target-revcomp(mCherry)-Donor, such that inversion yields mCherry expression. In the case of a split bRNA, the same construct was prepared, except with only one loop of the bRNA encoded. 50 ng each of two plasmids encoding one loop and the recombinase were cotransfected with the 288 ng of the same mCherry inversion reporter. In all cases, 1.4 x 10^4^ cells were transfected in 96-well plates with 0.5 µL well Lipofectamine 2000 per well. After 60 hours at 37°C, media was aspirated and cells were detached using 50µL of TrypLE Express (Gibco). TrypLE was neutralized with 50 µL Stain Buffer (BD), then, cells were resuspended then spun down in 96-well U-bottom plates at 300xg for 5 minutes. The supernatant was aspirated and the cells were resuspended in 120 µL Stain Buffer (BD). Flow cytometry was performed using the Quanteon Novocyte and resultant data was analyzed using NovoExpress software. Efficiency was recorded as the MFI of mCherry+ cells within the GFP+ gate or as the percentage of mCherry+ cells within the GFP+ gate.

### Genome insertion of plasmids

The activity of IS622 for genome insertion was measured in HEK293FT cells. For screening genomic sites, 137 ng of a plasmid encoding Ef1a-Rec-3xSV40NLS-P2A-EGFP was cotransfected with 60 ng of a plasmid encoding U6-bRNA and 534 ng of a plasmid encoding Ef1a-Puro^R^-P2A-mCherry and a recognition sequence for the bRNA. For assessing the efficiency and specificity of insertion with TBL or DBL binding the genome, 137 ng of a plasmid encoding U6-bRNA and Ef1a-Rec-3xSV40NLS-P2A-EGFP was cotransfected with 534 ng of a plasmid encoding Ef1a-Puro^R^-P2A-mCherry and a recognition sequence for the bRNA. For assessing the efficiency of insertion with engineered recombinases with various bRNAs, 137 ng of a plasmid encoding Ef1a-Rec-3xSV40NLS-P2A-EGFP was cotransfected with 60 ng of a plasmid encoding U6-bRNA and 534 ng of a plasmid encoding Ef1a-Puro^R^-P2A-mCherry and a recognition sequence for the bRNA. In the case of a split bRNA, 30 ng each of two plasmids, each one expressing one loop, were cotransfected. In all cases, 1.4 x 10^4^ cells were transfected in 96-well plates with 0.59 µL well Lipofectamine 2000 per well.

### Cell culture for selection of on- and off-target insertions

After 96 hours, all cells were passaged from 96-well to 24-well plates. After two additional days, 20% of cells were passaged in 10 µg/mL puromycin. Cells were passaged under puromycin selection for an additional 12 days, for a total of 18 days post-transfection.

### Mapping insertion sites genome-wide

After selection, genomic DNA was extracted from cells using *Quick*-DNA Miniprep Plus Kit (Zymo). Genomic DNA was measured with Qubit HS dsDNA Assay (Thermo). 150 ng of gDNA was then tagmented with Tn5 then amplified using a plasmid donor specific bait primer. Sample preparation, sequencing, and sequencing analysis for determination of insertion sites was performed as previously described (Durrant et al. 2022), with some modifications to account for a UMI of length 20 rather than of length 12. Samples were sequenced on an Illumina NextSeq with 600 cycle P1 or P2 kit.

### Selection of unique genomic sites for genome insertion

Genomic sites were selected from amongst 14 nucleotide sequences found only once within the hg38 reference genome. Of these, we filtered for sequences with either a CT or GT core sequence at position 8 and 9 of the 14 nucleotide sequence. Amongst these, we selected sequences that are between the transcriptional start site and 5 kbp downstream of the transcriptional start site. Sequences were further filtered by essentiality, using a published list of essential genes (Funk et al. 2022). Most sequences were then selected from amongst the top 30% of expression level in HEK293FT cells, based on RNA-seq data of unmodified cells (GEO: GSE164956) (Katrekar et al. 2022). Genome sequences were paired with plasmid sequences based on matching core sequences and a lack of a matching nucleotide at position 7, which interferes with the handshake base pairing mechanism (Hiraizumi et al. 2024). For intra-genomic inversion and excision, pairs of unique 14 nucleotide sequences were selected for relative distance to one another. Next, the pairs of sequences were filtered to require both to encode an identical core sequence. Finally, the sequences were filtered again to ensure position 6 and 7 of the sequences did not match.

### Selection of genome-orthogonal sequences

Sequences of length 14 that were orthogonal to the human genome were identified. Sequences were subsequently filtered for those containing a CT or GT at position 8 and 9 of the 14 nucleotide sequences. Next, sequences were compared against the genome to identify all genome locations matching positions 1-11 of the 14 nucleotide sequence. Genome orthogonal sequences were then rank-ordered according to the number of 11-mer off-targets in the genome, with zero being optimal.

### Generation of LSR landing pad cell line

A landing pad HDR template plasmid was designed to express the LSR Dn29 protein, with the attP sequence positioned between the Ef-1a promoter and the Dn29 coding sequence. This design enables a promoter swapping mechanism where delivery of a plasmid library containing the attB sequence interrupts Dn29 expression and activates expression of the library member. For selection purposes, a blasticidin resistance gene was linked to the Dn29 coding sequence via a P2A peptide, allowing for selection of successfully edited cells that maintain expression.

1 x 10^6^ K562 cells were nucleofected with the HDR template plasmid (5 μg) along with TALEN expression vectors targeting the AAVS1 locus: 1 μg each of hAAVS1 1L TALEN (Addgene #35431) and hAAVS1 1R TALEN (Addgene #35432). Nucleofection was performed using the Lonza SF Cell Line 4D-Nucleofector X Kit L according to the manufacturer’s protocol. Beginning three days post-nucleofection, the media was supplemented with 20 μg/mL blasticidin for 10 days, followed by maintenance at 10 μg/mL blasticidin. To establish monoclonal populations, single cells were sorted into 96-well plates using a BD FACSAria Fusion and expanded over a two-week culture period. Integration of the landing pad at the AAVS1 locus was validated by PCR. To evaluate Dn29 integration efficiency, cell lines were nucleofected with 1.2 μg of a promoterless plasmid containing the attB sequence upstream of an mCherry coding sequence. Eleven days after nucleofection, mCherry expression was quantified using an Attune NxT Flow Cytometer (Thermo Fisher), comparing results to wildtype K562 cells nucleofected with the same promoterless mCherry donor as a control for background expression.

### Design of libraries for IS622 recombinase deep mutational scan

To generate a comprehensive library of single amino acid substitutions in the IS622 recombinase, we ordered a 250 bp oligonucleotide pool from Twist Bioscience. This pool was divided into five sub-pools, each covering a distinct 65 amino acid stretch of the IS622 sequence: residues 2–66, 67–131, 132–196, 197–261, and 262–326. The library was designed such that each amino acid substitution was encoded by 2 different codons (when available) and was designed to include completely random 65 amino acid stretches that resulted in the formation of a full-length protein.

Each sub-pool was PCR-amplified in a 50 µL reaction using Kapa HiFi HotStart PCR Mix (Kapa Biosystems), with 10 ng of synthesized oligonucleotide as template and sub-pool-specific primers. The thermocycling parameters were: 98°C for 120 s; 10 cycles of 98°C for 10 s, 65°C for 15 s, and 72°C for 15 s; and 72°C for 1 min. PCRs were cleaned up using a 0.8x ratio of Agencourt AMPure XP beads (Beckman Coulter). Plasmid backbones for cloning each sub-pool were generated by replacing the corresponding sequence of the IS622 with BsaI recognition sites, to enable Golden Gate assembly of the libraries. Each backbone (3 µg) was digested with BsaI in a 50 µL reaction for 3 hours at 37°C, followed by heat inactivation at 65°C for 20 minutes. Digested products were purified using DNA Clean & Concentrator-5 columns (Zymo). Golden Gate assemblies were performed using 500 ng of backbone and 100 ng of insert split across multiple 50 µL reactions, incubated at 37°C for 3 hours and heat-inactivated at 65°C for 20 minutes. Reaction products were purified and eluted in 10 µL of water.

Electrocompetent Endura DUOs (Lucigen) were transformed with the assembled libraries following the manufacturer’s protocol. A small aliquot (1–5 µL) was plated on carbenicillin LB agar to estimate library coverage, while the remaining culture was expanded overnight in 70 mL LB with carbenicillin. Library coverage of at least 1000x was confirmed before downstream applications. Plasmid libraries were sequenced using the NextSeq 2000 (150 bp PE run).

### Generation of recombinase library cell lines

Each sub-library contained approximately 2,500 elements. Our K562-Dn29 landing pad cell line exhibited an integration efficiency of around 15%. To ensure a minimum of 1,000x coverage, we started with 2.5 x 10^7^ K562-Dn29 landing pad cells per replicate of the library and used 50 µg of library plasmid. Cells were divided into multiple 100 µL nucleocuvettes and nucleofected according to the manufacturer’s protocol (Lonza). Starting three days after nucleofection, the culture media was supplemented with 2 μg/mL puromycin for a minimum of seven days.

### Transfection and sorting of recombinase library cell lines

For each replicate, 3 x 10^7^ recombinase library cells were nucleofected with 60 µg of plasmid containing the bRNA and inverted mCherry reporter. Cells were divided into multiple 100 µL nucleocuvettes and nucleofected according to the manufacturer’s protocol (Lonza).Three days after nucleofection, 20% of the cells were collected as the pre-sort fraction. The remaining cells were sorted using a FACSAria Fusion Cell Sorter based on their mCherry to GFP fluorescence ratio. Post-sort, a minimum of 1,000x library coverage was maintained.

### Extraction and preparation of recombinase libraries

Genomic DNA (gDNA) was extracted using the Quick-DNA miniprep plus kit (Zymo), following the manufacturer’s protocol. To achieve a minimum of 1,000x coverage, at least 15 µg of gDNA was used for the first PCR (PCR1). For each replicate, 16 x 50 µL PCR1 reactions were prepared using Kapa HiFi HotStart PCR Mix (Kapa Biosystems) and primers that specifically amplify recombinases integrated into the genomic landing pad. PCR1 products were purified with a 0.7x volume of Agencourt AMPure XP beads (Beckman Coulter). The thermocycling parameters were: 98°C for 120 s; 24–27 cycles of 98°C for 10 s, 65°C for 15 s, and 72°C for 40 s; and 72°C for 2 min. For the second PCR (PCR2), 100 ng of purified PCR1 product was used to amplify recombinase regions corresponding to each library in a 50 µL reaction, again using Kapa HiFi HotStart PCR Mix. The thermocycling parameters were: 98°C for 120 s; 9-12 cycles of 98°C for 10 s, 65°C for 15 s, and 72°C for 15 s; and 72°C for 1 min. This PCR2 was cleaned up with 0.7x ratio of Agencourt AMPure XP beads. The third PCR (PCR3) was performed to add sample indices, using 100 ng of purified PCR2 product as input in a 50 µL reaction, using Kapa HiFi HotStart PCR Mix. The thermocycling parameters were: 98°C for 120 s; 5-7 cycles of 98°C for 10 s, 65°C for 15 s, and 72°C for 20 s; and 72°C for 1 min. PCR3 products were gel extracted using the Monarch DNA gel extraction kit (New England Biolabs) and then cleaned up with 0.7x ratio of Agencourt AMPure XP beads. In all the above PCRs, the numbers of cycles were tested to ensure that they fell within the linear phase of amplification. The libraries were quantified using the Qubit dsDNA HS assay kit (Thermo Fisher) and pooled together and sequenced on a 150 bp PE run on the NextSeq 2000 (Illumina).

### Analysis of deep mutational scan

Paired-end reads with overlapping regions were merged using BBMerge (Bushnell et al. 2017). To analyze the sequencing data, we developed a Python-based pipeline to count variant frequencies within specific regions. The script begins by initializing a dictionary from a reference library file containing all expected variant sequences. For each read in the FASTQ file, it searches for two constant flanking sequences within the merged read. If both flanking sequences are identified, the sequence between them is extracted and counted only if it matches exactly with an entry in the reference library.

For each sample, variant counts are normalized using the formula (count + 1) / total counts across all library members. Enrichment scores are calculated by dividing the normalized post-sort frequency by the pre-sort frequency for each variant. These enrichment scores are specific to each library. A z-score is then computed as z = (x - µ)/σ where x is the enrichment score of the variant, µ is the mean enrichment score of all members of a library, and σ is the standard deviation of the enrichment scores. The relative z-score is determined by subtracting the z-score of the library-specific wild-type (WT) variant from that of each individual variant.

Finally, mutability scores were calculated for each position by counting the number of variants whose relative z-score falls within one standard deviation of the WT.

### Prediction of structural contacts

To analyze the structural interactions of bridge recombinases with DNA and RNA substrates, we developed a computational approach for the identification of protein residues involved in critical contacts. We utilized the cryo-EM structure of IS621 (PDB: 8WT6) to predict and compare key interaction interfaces. Protein-nucleic acid and protein-protein contacts were identified using a distance-based approach implemented in Python (3.12.2) with Biopython (v1.84). Residues were classified as contacting if any atom in the residue was within 4.0 Å of any atom in a DNA, RNA, ion, or protein molecule. For each residue, we calculated the solvent-accessible surface area (SASA) using the Shrake-Rupley algorithm and classified residues as solvent-exposed (SASA ≥ 10.0 Å^2^) or internal. We further categorized protein contacts as monomer (intra-chain), dimer, or tetramer contacts based on the identity of the contacting chains.

### Modeling of recombinase mutations

Recombinase mutations were modeled using the IS621 transpososome structure (PDB: 8WT6) and modeled in ChimeraX. Amino acids were mutated using the swapaa command with Dunbrack rotamers. Hydrogen bonds for rotamers were predicted using the default settings, with a distance tolerance of 0.40 Å and an angle tolerance of 20°.

### ddPCR of genome insertion

Three days post-transfection, cells were trypsinized with 50 µL TrypLE (Gibco) for 10 minutes and then quenched with 50 µL Stain Buffer (BD). The 100 µL cell suspension was split into two 50 µL aliquots in U-bottom 96-well plates, centrifuged (300 × g, 5 minutes), and supernatant aspirated. One plate was resuspended in 120 µL Stain Buffer (BD) and analyzed with Novocyte Quanteon Flow Cytometer with Autosampler (Agilent). The other plate was resuspended in 50 µL QuickExtract DNA Solution (Biosearch Technologies), vortexed for 15 seconds, and thermocycled: 65°C for 15 minutes, 68°C for 15 minutes, 98°C for 10 minutes. DNA was cleaned with 0.9x AmpureXP (Beckman Coulter) beads. To assess integration efficiency, qPCR/ddPCR primers and probes were designed to span the left integration junction of various loci in the human genome, using a constant primer that binds to the donor plasmid sequence (PR_NPXX) a genome binding primer near the locus (PR_NPXX) and a FAM probe within the amplicon (pbNPXX).

Genomic reference primers and probes located nearby each insertion locus were designed to measure locus copy number for efficiency percentage calculations. ddPCR reaction mix (22 µL total): 11 µL ddPCR Supermix for Probes (no dUTP) (Bio-Rad), 1.98 µL of each primer (10 µM), 0.55 µL of each probe (10 µM), 1.65 µL cleaned gDNA, 0.22 µL SacI-HF (NEB), water to volume. Each reaction contained primers and probes for the target site (FAM probe) and a nearby reference locus (HEX probe). Reactions were run on QX200 AutoDG Droplet Digital PCR System (Biorad). For off-target detection or low concentration samples, primers were increased to 20 µM and volume halved, and gDNA volume was increased to 4.95 µL.

### Genomic inversion and excision and sample preparation

750 ng of a plasmid encoding U6-bRNA and Ef1a-Rec-P2A-EGFP was transfected into 1.4 x 10^4^4 HEK293FT cells using 0.59 µL Lipofectamine 2000 in 96-well plates. After 96 hours at 37°C, genomic DNA was extracted from cells using 50 µL of QuickExtract (Biosearch Technologies) per well. Extracted DNA was cleaned up using 0.9x AmpureXP magnetic beads (Beckman Coulter). 150 ng of genomic DNA was then tagmented with i5 handles using Tn5 transposase as previously described (Durrant et al. 2022). Tagmented gDNA was cleaned up using 0.9x AmpureXP magnetic beads. Then, a PCR was performed with a i5-specific primer and a second primer designed to bind upstream of the rearrangement junction. PCR was performed with 2x Platinum SuperFi with a protocol of 1 Cycle, 98°C for 2 min; 12 cycles, 98°C for 10 s, 68°C for 10 s, 72°C for 90 s; 1 cycle 72°C for 5 min. After cleanup with magnetic beads, a second PCR was performed with an outer nested i5 primer and a full-length i7 primer with a binding region specific for a region downstream of the first primer but upstream of the rearrangement junction. PCR was performed with 2x Platinum SuperFi with a protocol of 1 Cycle, 98°C for 2 min; 22 cycles, 98°C for 10 s, 68°C for 10 s, 72°C for 90 s; 1 cycle 72°C for 5 min. Genome specific primers were designed using Primer3. For both inversion and excision, genome specific primers were designed such that they bound upstream of a junction but not within the region being inverted or excised. Samples were run on a 2% agarose gel and then gel extracted from 200-600 bp using Monarch DNA Gel Extraction Kit (NEB). Samples were then sequenced on an Illumina NextSeq with 600 cycle P1 or P2 kit.

### Measurement of inversion and excision efficiency

Fastq files of sequenced amplicons were trimmed and assembled using FLASH and Cutadapt. Human genome reference sequences were prepared for the measured junction. One fasta file was prepared with the wild-type genom sequence, and a second fasta file with the expected rearrangement product based on the sequence pair. Assembled reads were separately mapped to both files using bwa. Alignments were then filtered, requiring reads to overlap with 10nt upstream of the core sequence and 20 nt downstream of the core sequence. The resultant alignments contain only reads that align exactly the expected rearrangement junction. The number of reads in both files was counted, and the rearrangement efficiency was calculated as the number of reads in the rearranged alignment divided by the number of reads in both the wild-type and rearranged alignments.

### Excision of *BCL11A* enhancer sequences

The *BCL11A* DHS58+ sequence was cloned into a plasmid such that the regions upstream and downstream of the GATA motif were separated by a 1 kbp sequence containing a terminator. This cassette was between a Ef1a promoter and mCherry gene, such that recombination between the two regions of *BCL11A* would result in excision and mCherry expression. 100 ng of a plasmid expressing a U6-bRNA and Ef1a-Rec-P2A-EGFP was cotransfected with 372 ng of this reporter plasmid into 1.4 x 10^4^ HEK293FT cells using 0.5 µL Lipofectamine 2000 in 96-well plates. After 60 hours at 37°C, media was aspirated and cells were detached using 50 µL of TrypLE Express (Gibco). TrypLE was neutralized with 50 µL Stain Buffer (BD), then, cells were resuspended then spun down in 96-well U-bottom plates at 300xg for 5 minutes. The supernatant was aspirated and the cells were resuspended in 120 µL Stain Buffer (BD). Flow cytometry was performed using the Quanteon Novocyte and resultant data was analyzed using NovoExpress software. Efficiency was recorded as the MFI of mCherry+ cells within the GFP+ gate or as the percentage of mCherry+ cells within the GFP+ gate.

### Excision of GAA repeats

GAA repeats sequences were cloned into a plasmid such that two GAA regions were separated by 1 kbp of intervening sequence, such that recombination between the two GAA regions results in excision. 100 ng of a plasmid expressing a U6-bRNA or U6-DBL and Ef1a-Rec-P2A-EGFP was cotransfected with 372 ng of this reporter plasmid into 1.4 x 10^4^ HEK293FT cells using 0.5 µL Lipofectamine 2000 in 96-well plates. After 72 hours at 37°C, genomic DNA was extracted from cells using 50 µL of QuickExtract (Biosearch Technologies) per well. Extracted DNA was cleaned up using 0.9x AmpureXP magnetic beads (Beckman Coulter). Primers were designed such that amplification would result in equal amplification of the unmodified and modified plasmids. PCR was performed with 2x Platinum SuperFi with a protocol of 1 Cycle, 98°C for 1 min; 20 cycles, 98°C for 10 s, 68°C for 10 s, 72°C for 10 s; 1 cycle 72°C for 2 min. PCR products were run on a 2% agarose gel, stained with SYBR Gold and visualized with a BioRad ChemiDoc Imaging System. The lower band was then gel extracted and sequenced using the Premium PCR service from Plasmidsaurus. The number of repeats between the primer binding regions was counted using custom python scripts. Then, the count table was filtered for repeat counts possible via intra-plasmid excision rather than via inter-plasmid recombination. The number of repeats excised was calculated as the number of total repeats in the initial plasmid minus the number of repeats counted.

## Data Availability

Data supporting the results are in the main text, figures, and supplementary tables. Next generation sequencing datasets generated as part of this study will be available on the NCBI Sequence Read Archive prior to publication. Relevant plasmids will be made available at Addgene.

## Acknowledgments

We thank all members of the Hsu lab for helpful input throughout the course of the project. We are grateful to Januka Athukoralage for insights on the bridge recombination mechanism, and Vincent Tran for discussions about deep mutational scanning. We also thank Brian Plosky, Rachel Senturia, and Joseph Caputo of the Arc Institute. N.T.P., L.J.B., D.K., G.A.G., M.G.D., J.J.P., A.F., and S.K. are supported by funding from the Arc Institute. M.H. is supported by JSPS KAKENHI Grant Number 23K14133, Takeda Medical Research Foundation, and JST, ACT-X Grant Number JPMJAX232F. H.N. is supported by JSPS KAKENHI Grant Numbers 21H05281, 22H00403, and 25H00436, Takeda Medical Research Foundation, the Inamori Research Institute for Science, and JST CREST Grant Number JPMJCR23B6. P.D.H. is supported by funding from the Arc Institute, Rainwater Foundation, Curci Foundation, Rose Hill Innovators Program, S. Altman, V. and N. Khosla, and anonymous gifts to the Hsu Lab.

## Author contributions

N.T.P, M.G.D., S.K., and P.D.H. conceived the study. N.T.P., L.J.B., D.K., S.K., and P.D.H. designed experiments. N.T.P., L.J.B., D.K., G.A.G., and J.J.P. performed experiments. N.T.P., L.J.B., D.K., G.A.G., M.G.D., M.H., H.N., S.K., and P.D.H. analyzed and interpreted data. D.K. designed deep mutational scan strategy. A.F. designed and prepared engineered cell lines. N.T.P. and M.G.D. performed off-target analyses. M.G.D. nominated orthologs for testing. G.A.G. analyzed AlphaFold structures. M.H. and H.N. advised on bRNA and protein engineering. N.T.P. and C.R.T. designed figures with assistance from D.K. N.T.P. and P.D.H. wrote the manuscript with important contributions from H.N. and S.K. and input from all other authors.

